## Supplementary material for "Octopamine and tyramine signaling in *Aedes aegypti:* characterization, distribution and potential role in the Dengue vector development and physiology": S1. AaOAa1-R.docx

**Supplementary figures 1.** *Aedes aegypti* α1 octopamine receptor (AaOAα1-R)

**Figure S1a.**

TGCCGTTTGGTTGTAAAATTGCGAATCGTTCGCCGAGACGCGGGACAAGTGCTCTGCACGGTTGTTATCAGTCTCTACTAACTAGTCCTAGTAGCTAGCTGGGATGCTTCCATCATCATCATCATCATCATCATCGTCATCAGTGTATAGACCTTCTGCCAGTGGGGCTGCCCTGGGAGGAAAGAGTGTAACGCTCGCTTCCGGTTTCCGTGTCCGCCCGTCAGGTTGAAAGAGTGAAGTTCTTCAGTGTAGACGCGAAGTGGTTTCCCGACGGAACTAGTGTGGATTTTTTCGGGGGGTTATTCCAAGCAGAAAGTCGCAGTGTGCCTAGAGATAATATTTCGACTCCGTCTTTTGAAGCGAAATGCCCTGGCAGAATTTGGCCTGCTGTGATGCGCACATCAATCAGCCTACTGTAGAAATGCTAGGTGTAATGTAAGCTCACTCTGTCTACGTGCTGTGGGTGACGAGTTAATAGCAAAGTACATGAAACCAAAGTGCTAAAATAAGCGTGAAATGGATTTTCGACGAGAAACCGGTGAGGAAATCTGATAGTGTGTCTCGTTTGTGCCCGTTAGTAGTGCTGGCTGCTGTGCTGTATTGTTACCATCGAGAAGCTCCACAGCGATTGTGATGGTGCGGGTTTACGGTTTGTTTACGGCTGTGGAAGTTTAGCGCCTGAAACGGAAGGCAGGAAGGCAGAATACGGAAGGAGAAGAAACCAAGCTAGAAGCCTTTAAAAAGTTGCATCTCGGTAACTCGGTGTTTCCCTTGGCGAACCGTCGAAGCAGGCTCTGACGCTGCTAGTAGTTCGACTTCTACCGCGACGTGAGTGTGCATTGAGAAAAGAAATAAAAGTATTTAGACGTAAGCTGATAAGATTCCTGGGGTGAATTACAGAAAAAAAGTGTTGTGGCTTCACGGTTTGAGCAGTGGTTTTGTTAATCGAAGGAGAAAAGCGCATCCAGAAATTCCAGTAGCTTGCAGTCAGTGTTTGTTCGACAGCAAGGAAAACTTGTTCCAGCAATTGCCGCTTGGCTGCGTGGAGTCCGAAAACAAATTGTGATTATCGCAGCAGGCGTGGGCTGATGTAAATCTCCAGTTCGAAATTGGAGACTGTTGTTGCTAATGTTCGCGACTTGAAAGCTGATCGGCGAGGACTTGCCGTCGTCGACCATGAAAGAAATGTGCAATCCGCAGATCAAACGGAATCATTGAAAAGAATCCTTGAGAACGTGGTGAATCGAGTGCAGTGAAGTGAAGACGACCCCATAAATACACAAACACACACACACACATACACAGAGAAGGAGAGATAGACAGAGATAGAAGCCATCATCCGTTCACAGTTCACAGTGTACGGTGACAACAAATTGAAATGTGAAAACGGCAAGTTGGAAATCGAAAAGCACTTTAGAGCCGAACAGAGAACCGGAGGAAAAACCGGAAGGCGCCCATTTCTCTTTCAATAGCAAGCAATTTCCAGACCATTAGTGGGGGCATTTGTGACAGCTATACAAAGCGAGTGAGAGAGAGAGAGCTAGCATCCATCCTTGAAGTTTTGAACTGCTCGTGCAAAGGTTGCTTTGTTGGTTGCTGTCGATGTGTTCTTGAATTGTGGAGGGGGTTTCTGAGGTCAATGTTGAGGAAAGGAAATCAAGTGGAACGAGACCAAGAGCGCAAATCAAGGGAAAGCCAGAAGTAGTTGCATTAGCGAAGAAGTTCAAGGTGGGGGTAACATCTTTCCATTCTGCTGCTCTCCAAGGGTGGGGGCAAGAAGTGTTCGGTTTTGGTGTTGCTCCGGAGGCGCTTGTGTGTCGGTTGTGTGGGTTGAATGGCTACTACCGGACGACATCGACAGCAACGTCAACGAGGTGCAGCAGATAGAAGCAATAGCAGAAGCAGATTCAGCAGCATCAACATCAACAGCAACACCGCCGTGAAGAAGAGTGCTGTCAGTGTCAACGGTTTGTGTTCGTCATCGTCGGAATCGTTGGCCGTTGGCGTGTTACGTGAACCGCAAAATGGCTAGATTCTGCCTGCGATTCTGGTCCGAGCTGAGATTTGTTAGAGCAAACGAGAGTTTCACTTCTACTACCGGAGTCGACTGTAGCACAGCGACATTCCACACCACAGAAGCCGGCCCTGTCGGCAAGAAATCTACAATCAGAAGAAAGAGATACAGAAGAGTGGCCTTTGTTGGTGGCGTTCGGTTATCGGCAAGTCGTTCGGATCGAAGCAAGCTGCCATCGCTGTCACTCTGCAACAGAGTTGGAGATGATCACCGGCCACCACTAGGACCGGAAGAGTCGGCAGTGGTTGGTCATCGCTTACCGTTGTGCGCCGGAATCGAACCGCCGCGGAGCAGCGTCCGAG**ATG**AATGCGACCGAGTGCAACAGCCTGATTGCCTCGGTCGAGTGGACCGAGCCGCGCAAGATCCTGCTGCTGGCCATCCTCATCTTCATCGACATTTTGGTCATCGTCGGGAACACGCTCGTGGTGGCCGCCGTCACCAGTTCGCATAAGCTTAGGAGTGTTACCAATTTTTTCATCGTTAGTCTAGCCGTTGCCGATTTGTTAGTTGGATTAGCTGTATTGCCGTTTTCGGCCACCTGGGAGGTGTTTAAGGTATGGATCTTCGGTGACGTGTGGTGCCGCGTTTGGCTGGCCGTCGATGTCTGGATGTGCACCGCTTCGATACTGAACCTGTGCGCGATTTCGCTCGACCGCTACGTGGCCGTCACCCGTCCCGTCACCTACCCGAGCATCATGTCCACCAGGAGGGCCAAGTCGCTCATCGCCGGGCTGTGGGTGTTGTCCTTTGTGATCTGCTTTCCGCCGCTGGTCGGGTGGAAGGAGCAGAAAGTGAAAGAGAACCTATTATTACCGTATGGAAATCACACGCTGGTGTCAATGTCACCGTCACCAGTCACTTCCACCTCATCGTATTCATCATCGCCATCAGTGTCGTCGATATCATCATCGTCTTCATCGCCATCAACAGGAACGTCCTCGGCTTTCAGCACCATAACCTCCTCCTCGATTGCCGAGAGCTGGGAAGTGGGAGGGATGGGCGAAACGTCCCCCAATCCGCCCCCCTGCCCGTGGACCTGCGAGCTGACAAACGACGCCGGATACGTCGTCTACTCGGCCCTGGGGTCCTTCTACATCCCCATGTTCGTGATGCTGTTCTTCTACTGGCGCATTTATCGGGCCGCCGAGCGGACCACACGTGCCATCAATCAGGGATTTAGGACCACCAAAGGTATGGGCACCCGTTTCGATGACAACCGGCTGACGTTGCAGATTCATCGTGGCCGCGGTTCGACGGCCAGTGCCCACGGAGCGCCCACCATTGGTGGCAGTGCTGCCGGAGTTCTTGGCAGTGCCCATCACGAGTCCCCGCACAGCAACGGAAGCACCCACAGTACTACCACCAGCATTGGCAGTGCCTCGCCGGAACGCCTTTCGCGATACATGACCCGCTGCAAGAACCACGAGAAGATCAAAATTTCCGTGTCCTACCCGTCGACGGAGAATCTGAACCAGAGCTCCTCGGCTGGAGAAGGCAGTAAGCTGCTGTACGCGGTGCACTATTCTTCGAACGGAGGCCGAGAACACAACGCTTCGCACATCTTCCGAAGGCCTTCCAAGGAGCAGAGTGGCAGCGGTCAGTATCTAACGGTGGACGGCCAAGGAAGAAGCTTGCTGTCGCCGCGATCCAGCAAACGAATGGGCAAGCGTAACATCAAAGCTCAGGTCAAGCGCTTCCGGATGGAAACCAAGGCGGCCAAAACGTTGGCCATCATCGTCGGGTTGTTCATCCTGTGTTGGCTTCCGTTCTTTACCATGTACCTGATAAGGCCCTTCTGTGACAACTGCATCAACGATCTGCTGTTTTCGATCGTGTTCTGGATCGGGTACTGCAACTCGGCCATCAATCCGATGATCTACGCCCTGTTTTCCAAGGACTTCCGGTTCGCATTCAAGCGGCTGATCTGCCGATGTTTCTGCTCGGCTGAGGCCATTCCAAGGCCGGCCAGTCGGCGAGGATCGGACATGTCGCAGATACGGATGCACGGGGCGCGAACTCCGAGCATATCGCCATCGGCGGCAGCGCAATCCATTGGCGACGACAGCGACCCGGTGGGAGACCTGTCCGACAGCAGA**TGA**CGATCGTTTGGCCCGAGCGTCAGTTCGAATTCTAGTCCGGGCCTGGGCAGCGGAACACGTCGTCGCCGTAGCAACAGTATTACCAATGCTGCATCGTCGGCCGGTGGAACCATGGCAACGGCAACGACGATGAGCAGCACGTCGCCTACTGTGGACGCGTCCGGAACTAGGGGCATCGTCAGTCGGGCGCCGGTCTACGACATCTGCCATCCCCTGGTAATGGAAAGCCCGGGGTTCACCGGGTACGGGAGCAGCGCCGGCGGTAGGTTCGGGGGCGAGGATGGACTGACGCGGGGGTCAGACAGTGCGAGGGAAGGAAGCGTCGGCGGCAGCAGCTTGACCAGGGTGGATATTTGATCGTTCTGGGGCAGGAGATTGCTTCACTTCTGTGACAGCTCGTAAGTTTGGTTCGATGTGACGGGGACGATTATTCCGCGTGGTTTTAAGGAGGTTCGGGCGCATAGAATACCTTGGTATCCACAACTAATTTGAAACTTTTTTTAAGATCTTCGACGATCAGCTCGATCGCAGTGGCACTGCGTTCACAAAGTTTCCATGATAATGACTGTGGTGAATATACCTTATCTAGTTGTCTTTACTCACCACGCGGCGTTAAGCCCCAAACACAAAGTAAACGGCAGCACAGTCCAACGGCAGAACGGCAAACCCTACTTGATCCACGGTTAACTTATCAGGCCTATGTTGAATCTTGTATAGACGACACTGAACAAAACATTGACATGTCATTCAAATCGATGT

**Figure S1a.** *Aedes aegypti* α1 octopamine receptor (AaOAα1-R) complete annotated sequence (XM_021839348.1). The primer used to amplify AaTAR1 were highlighted in blue. The start and the stop codon were highlighted in yellow.

**Figure S1b.**

ATG AAT GCC ACC GAG TGC AAC AGC CTG ATT GCC TCG GTC GAG TGG ACC GAG CCG CGC AAG ATC CTG CTG CTG GCC ATC CTC ATC TTC ATC GAC ATT TTG GTC ATC GTC GGG AAC ACG CTC GTG GTG GCC GCC GTC ACC AGT TCG CAT AAG CTT AGG AGT GTT ACC AAT TTT TTC ATC GTT AGT CTA GCC GTT GCC GAT TTG TTA GTT GGA TTA GCT GTA TTG CCG TTT TCG GCC ACC TGG GAG GTG TTT AAG GTA TGG ATC TTC GGT GAC GTG TGG TGC CGC GTT TGG CTG GCC GTC GAT GTC TGG ATG TGC ACC GCT TCG ATA CTG AAT CTG TGC GCG ATT TCA CTC GAC CGC TAC GTG GCC GTC ACC CGT CCC GTC ACC TAC CCG AGC ATC ATG TCC ACC AGG AGG GCC AAG TCG CTC ATC GCC GGG CTG TGG GTG TTG TCC TTT GTG ATC TGC TTT CCG CCG CTG GTC GGG TGG AAG GAG CAG AAA GTG AAA GAG AAC CTA TTA TTA CCG TAT GGA AAT CAC ACG CTG GTG TCA ATG TCA CCG TCA CCA GTC ACT TCC ACC TCA TCG TAT TCA TCA TCG CCA TCA GTG TCG TCG ATA TCA TCA TCG TCT TCA TCG CCA TCA ACA GGA ACG TCC TCG GCT TTC AGC ACC ATA ACC TCC TCC TCG ATT GCC GAG AGC TGG GAA GTG GGA GGG ATG GGC GAA ACG TCC CCC AAT CCG CCC CCC TGC CCG TGG ACC TGC GAG CTG ACA AAC GAC GCC GGA TAC GTC GTC TAC TCG GCC CTG GGG TCC TTC TAC ATC CCC ATG TTC GTG ATG CTG TTC TTC TAC TGG CGC ATT TAT CGG GCC GCC GAG CGG ACC ACA CGT GCC ATC AAT CAG GGA TTT AGG ACC ACC AAA GGT ATG GGC ACC CGT TTC GAT GAC AAC CGG CTG ACG TTG CAG ATT CAT CGA GGC CGG GGT TCG ACG GCC AGT GCC CAC GGA GCG CCC ACC ATT GGT GGC AGT GCT GCC GGA GTT CTT GGC AGT GCC CAT CAC GAG TCC CCG CAC AGC AAT GGA AGC ACG CAC AGT ACT ACC ACC AGC ATT GGC AGT GCC TCG CCG GAA CGC CTT TCG CGA TAC ATG ACC CGC TGC AAG AAT CAC GAG AAG ATC AAA ATT TCG GTC TCC TAC CCG TCG ACG GAG AAT CTG AAC CAG AGC TCC TCG GCT GGA GAA GGC AGT AAG CTG CTG TAC GCG GTG CAC TAT TCT TCG AAC GGA GGC CGA GAA CAC AAC GCT TCG CAC ATC TTC CGA AGG CCT TCC AAG GAG CAG AGT GGC AGC GGT CAG TAT CTT ACG GTG GAC GGC CAA GGT AGA AGC TTG CTG TCG CCG CGA TCC AGC AAA CGA ATG GGC AAG CGT AAC ATC AAA GCT CAG GTC AAG CGC TTC CGG ATG GAA ACC AAG GCG GCC AAA ACG TTG GCC ATC ATC GTC GGG TTG TTC ATC CTG TGT TGG CTT CCG TTC TTT ACC ATG TAC CTG ATA AGG CCC TTC TGT GAC AAC TGC ATC AAC GAT CTG CTG TTT TCG ATC GTG TTC TGG ATC GGG TAC TGC AAC TCG GCC ATC AAT CCG ATG ATC TAC GCC CTG TTT TCC AAG GAC TTC CGG TTC GCA TTC AAG CGG CTG ATC TGC CGA TGT TTC TGC TCG GCT GAG GCC ATT CCA AGG CCG GCC AGT CGG CGA GGG TCG GAC ATG TCG CAG ATA CGG ATG CAC GGG GCG CGA ACT CCG AGC ATA TCG CCA TCG GCG GCA GCG CAA TCC ATT GGC GAC GAC AGC GAC CCC GTG GGA GAC CTG TCC GAC AGC AGA TGA

MNATECNSLIASVEWTEPRKILLLAILIFIDILVIVGNTLVVAAVTSSHKLRSVTNFFIVSLAVADLLVGLAVLPFSATWEVFKVWIFGDVWCRVWLAVDVWMCTASILNLCAISLDRYVAVTRPVTYPSIMSTRRAKSLIAGLWVLSFVICFPPLVGWKEQKVKENLLLPYGNHTLVSMSPSPVTSTSSYSSSPSVSSISSSSSSPSTGTSSAFSTITSSSIAESWEVGGMGETSPNPPPCPWTCELTNDAGYVVYSALGSFYIPMFVMLFFYWRIYRAAERTTRAINQGFRTTKGMGTRFDDNRLTLQIHRGRGSTASAHGAPTIGGSAAGVLGSAHHESPHSNGSTHSTTTSIGSASPERLSRYMTRCKNHEKIKISVSYPSTENLNQSSSAGEGSKLLYAVHYSSNGGREHNASHIFRRPSKEQSGSGQYLTVDGQGRSLLSPRSSKRMGKRNIKAQVKRFRMETKAAKTLAIIVGLFILCWLPFFTMYLIRPFCDNCINDLLFSIVFWIGYCNSAINPMIYALFSKDFRFAFKRLICRCFCSAEAIPRPASRRGSDMSQIRMHGARTPSISPSAAAQSIGDDSDPVGDLSDSR*

**Figure S1b.** Nucleotide sequence of the α1 octopamine receptor (AaOAα1-R) open reading frame cloned from *Aedes aegypti* and deduced amino acid sequence. The primer used in RT-qPCR analysis were highlighted in red.

**Figure S1c.**

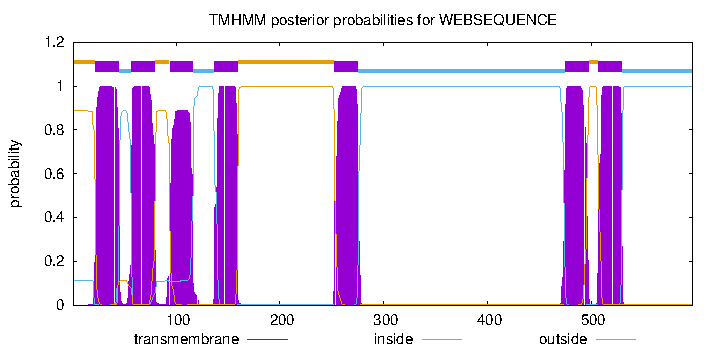

**Figure S1c.** Prediction of the AaOAα1-R transmembrane segments obtained with TMHMM v. 2.0 software.

**Figure S1d.**

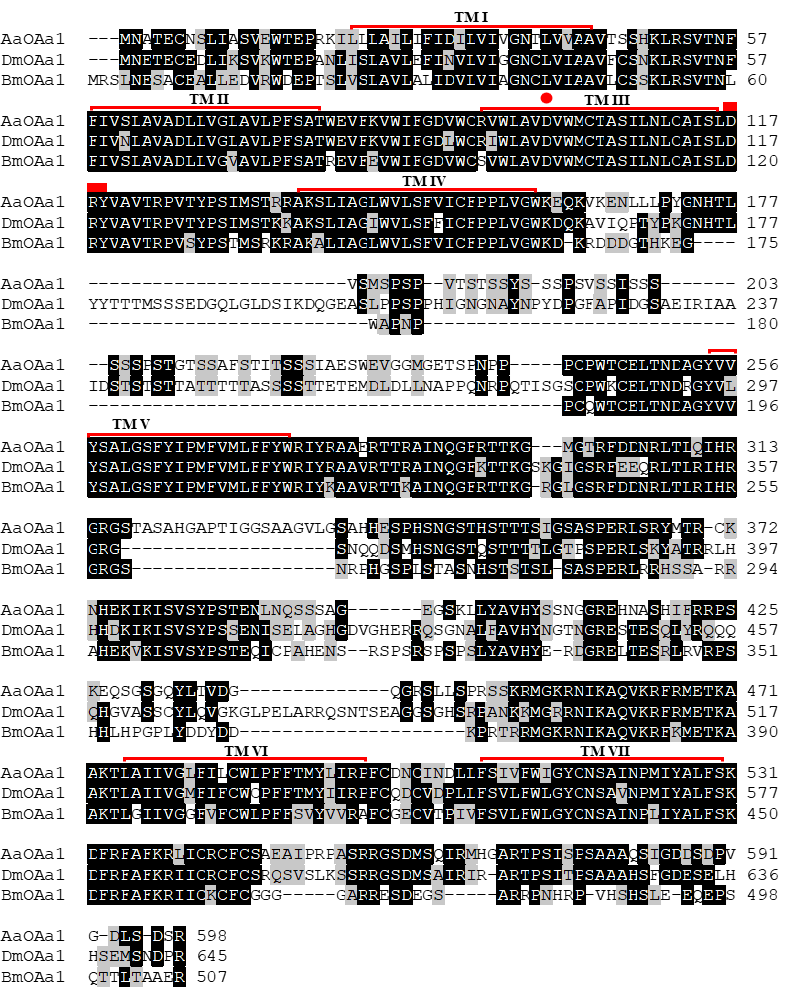

**Figure S1d.** Amino acid sequence alignment of AaOAα1-R with orthologous receptors from *D. melanogaster* (DmOAα1-R) and *B. mori* (BmOAα1-R). The putative seven transmembrane domains (TM I–VII) are indicated by red lines. Identical residues are highlighted in black while conservative substitutions are in grey. A red dot indicates the conserved aspartic acid and the serine residues that could interact with TA.
