## Supplementary material for "Octopamine and tyramine signaling in *Aedes aegypti:* characterization, distribution and potential role in the Dengue vector development and physiology": S2. AaOAa2-R.docx

**Supplementary figures 2.** *Aedes aegypti* α2 octopamine receptor (AaOAα2-R).

**Figure S2a.**

**ATG**GATTACCCAATGATAGCAACGGGCAACATCTCGCAGGAAGACTTCCTCGCCATAGTCGGGGCGAACTCTTCGCTCGCCAGCTTAATCCACGGATCGGCAACGACCGCCCTAAACGACTCCCTGGGCGGAGTGATAGCAACCGGCGACAACCGAACACTCTTCCACGGCTACCCGAGCGGATACACACTGCCGCACATCATCATCGCTTCAATTATCGTGACAATACTAATGATAATCATCGTAGTCGGTAACATGCTGGTGATCATAGCGATTGCCACCGAAAAGGCGCTCAAGAACATCCAGAACTGGTTCATAGCATCGCTGGCGGTGGCGGACTTCTTTCTGGGTCTGGTGATCATGCCCTTCTCGTTGGCCAACGAGCTGATGGGCTACTGGATCTTCGGGAACTGGTGGTGCGACATCCACTCGGCGATGGATGTGTTGTTGTGTACTTCGTCGATCATGAATCTCTGCCTGATCTCGCTGGATCGGTACTGGAGCATAACGAAGGCGATCGAGTATCTCAAATCGAGGACTCCGGGCCGGGCGGCTTTTATGATCGCTGCCGTGTGGATCATGTCGGCACTGGTGTGCATACCGCCGCTGCTGGGCTGGAAGGCGCCACGTCCGGAGGAGCACGTCGAACTTCCACAATGTCAGTTGAGTCAAGACATTGGCTACGTGCTGTACTCGGCGCTCGGATCGTTCTACATTCCCAGCTGCATCATGGTGTTCGTCTACATTAGGATCTACTACGCGGCAAAGGCCAGGGCCCGGCGGGGGATCCGGAAGAAGCCGCTCAAACCGCCCAGCGAGCAGGACACAAGCTTCACGCGTCCAGCGAATCCGATGCCATCGCTTCCCAGTGCCTCATCGATGTCGGCCGTTGCGGTGGCTCCGGCGTCATCCGTCAACGGAAACGGCCAATACGCCACGGCCAACAGCAACAACAACAATAACCTAAGTAGTGGCCCCAACAATCCTGGCCAGCATCAGCAGCAGATAGCGACGATAGAACCTCCACCGAGGGGGCTGGCGGGAGGTGCAGCTGGTGGTCCAACGCAGCAGATGTCGATCCCGACGGTGACGTGCGACTTCGCATCCGATATGTCCACCAGCGAAGCAGCCGACCATGACCATGCCGCTGTGGTAGCGGAGATGAAGGATACGCTCAAAGTCGGTAAACCGGGGATCGGGCTAACGTCACTGCCATTGCTGAGAGGGAACAACGGTTCGGCGGCCAGCAGTTCGGGGGCGGTGATTAGTCCGGGGTTAGCTAGGAATAGAGCACTTTCGGTAGGGATCGAAACCGATATGGTTAGCGAAATTGATCCATCGAGTTCCGACTCGGGGGTGATAAGTAGGTGTGCGGTGGTGAAACCGCTCAAGTTTAGGCTGTGTCAACCAATTTTCGGCAAGAGAGTAGGTAAGGGTCGTAATAAGAACAATAATGCGAGGGGAAATTGCGCAGGGGTGGGACCGGCAGCGATGGGAGGTCAAGCGATATCGATATCGGCCAAACAGAACGATAGCGCTTGTAGCGAACTGGAACCTGCACTGCCAAAGCCGAAGCCACGGGACCCGGAGAAAGAAAAGCGACGGATAGCGCGGAAGAAGGAGAAGCGAGCTACACTGATTCTGGGGTTAATTATGGGAAGTTTTATTGCCTGCTGGTTGCCATTTTTCTTCCTTTACATATTGGTGCCCGTTTGCAAGGACTGTCACATACCGGACTTTGCTTTCTCGTTGGCCTTTTGGTTGGGTTACATGAACTCGACTCTGAACCCGGCAATCTACACCATCTTCAACAAGGACTTCCGGAGAGCCTTCAGGCGGATCCTGTTCAAG**TGA**

**Figure S2a.** *Aedes aegypti* α2 octopamine receptor (AaOAα2-R) complete annotated sequence (KC439536.1). The primer used to amplify AaOAα2-R were highlighted in blue. The start and the stop codon were highlighted in yellow.

**Figure S2b.**

ATG GAT TAC CCA ATG ATA GCA ACG GGC AAC ATC TCG CAG GAA GAC TTC CTC GCC ATA GTC GGG GCG AAC TCT TCG CTC GCC AGC TTA ATC CAC GGA TCG GCA ACG ACC GCC CTA AAC GAC TCC CTG GGC GGA GTG ATA GCA ACC GGC GAC AAC CGA ACA CTC TTC CAC GGC TAC CCG AGC GGA TAC ACA CTG CCG CAC ATC ATC ATC GCT TCA ATT ATC GTG ACA ATA CTA ATG ATA ATC ATC GTA GTC GGT AAC ATG CTG GTG ATC ATA GCG ATT GCC ACC GAA AAG GCG CTC AAG AAC ATC CAG AAC TGG TTC ATA GCA TCG CTG GCG GTG GCG GAC TTC TTT CTG GGT CTG GTG ATC ATG CCC TTC TCG TTG GCC AAC GAG CTG ATG GGC TAC TGG ATC TTC GGG AAC TGG TGG TGC GAC ATC CAC TCG GCG ATG GAT GTG TTG TTG TGT ACT TCG TCG ATC ATG AAT CTC TGC CTG ATC TCG CTG GAT CGG TAC TGG AGC ATA ACG AAG GCG ATC GAG TAT CTC AAA TCG AGG ACT CCG GGC CGG GCG GCT TTT ATG ATC GCT GCC GTG TGG ATC ATG TCG GCA CTG GTG TGC ATA CCG CCG CTG CTG GGC TGG AAA GCG CCA CGT CCG GAG GAG CAC GTC GAA CTT CCA CAA TGT CAG TTG AGT CAA GAC ATT GGC TAC GTG CTG TAC TCG GCG CTC GGA TCG TTC TAC ATT CCC AGC TGC ATC ATG GTG TTC GTC TAC ATT AGG ATC TAC TAC GCG GCA AAA GCC AGG GCC CGG CGG GGG ATC CGG AAG AAG CCG CTC AAA CCG CCC AGC GAG CAG GAC ACA AGC TTC ACG CGT CCA GCG AAT CCG ATG CCA TCG CTT CCC AGT GCC TCA TCC ATG TCG GCC GTT GCG GTG GCT CCA GCG TCA TCC GTC AAC GGA AAC GGC CAA TAC GCC ACG GCC AAC AGC AAC AAC AAC AAT AAC CTA AGT AGT GGC CCC AAC AAT CCT GGC CAG CAT CAG CAG CAG ATA GCG ACG ATA GAA CCT CCA CCG AGG GGG CTG GCG GGA GGT GCA GCT GGT GGT CCA ACG CAG CAG ATG TCG ATC CCG ACG GTG ACG TGC GAC TTC GCA TCC GAT ATG TCC ACC AGC GAA GCA GCC GAC CAT GAC CAT GCC GCT GTG GTA GCG GAG ATG AAG GAT ACA CTC AAA GTC GGT AAA CCG GGG ATC GGG CTA ACG TCA CTG CCA TTG CTG AGA GGG AAC AAC GGT TCG GCG GCC AGC AGT TCG GGG GCG GTG ATT AGT CCG GGG TTA GCT AGG AAT AGA GCA CTT TCG GTA GGG ATC GAA ACC GAT ATG GTT AGC GAA ATT GAT CCA TCG AGT TCC GAC TCG GGG GTG ATA AGT AGG TGT GCG GTG GTG AAA CCG CTC AAG TTT AGG CTG TGT CAA CCA ATT TTC GGC AAG AGA GTA GGT AAG GGT CGT AAT AAG AAC AAT AAT GCG AGG GGA AAT TGC GCA GGG GTG GGA CCG GCA GCG ATG GGA GGT CAA GCG ATA TCG ATA TCG GCC AAA CAG AAC GAT AGC GCT TGT AGC GAA CTG GAA CCT GCA CTG CCA AAG CCG AAG CCA CGG GAC CCG GAG AAA GAA AAG CGA CGG ATA GCG CGG AAG AAG GAG AAG CGA GCT ACA CTG ATT CTG GGG TTA ATT ATG GGA AGT TTT ATT GCC TGC TGG TTG CCA TTT TTC TTC CTT TAC ATA TTG GTG CCC GTT TGC AAG GAC TGT CAC ATA CCG GAC TTT GCT TTC TCG TTG GCC TTT TGG TTG GGT TAC ATG AAC TCG ACT CTG AAC CCG GCA ATC TAC ACC ATC TTC AAC AAG GAC TTC CGG AGA GCC TTC AGG CGG ATC CTG TTC AAG TGA

MDYPMIATGNISQEDFLAIVGANSSLASLIHGSATTALNDSLGGVIATGDNRTLFHGYPSGYTLPHIIIASIIVTILMIIIVVGNMLVIIAIATEKALKNIQNWFIASLAVADFFLGLVIMPFSLANELMGYWIFGNWWCDIHSAMDVLLCTSSIMNLCLISLDRYWSITKAIEYLKSRTPGRAAFMIAAVWIMSALVCIPPLLGWKAPRPEEHVELPQCQLSQDIGYVLYSALGSFYIPSCIMVFVYIRIYYAAKARARRGIRKKPLKPPSEQDTSFTRPANPMPSLPSASSMSAVAVAPASSVNGNGQYATANSNNNNNLSSGPNNPGQHQQQIATIEPPPRGLAGGAAGGPTQQMSIPTVTCDFASDMSTSEAADHDHAAVVAEMKDTLKVGKPGIGLTSLPLLRGNNGSAASSSGAVISPGLARNRALSVGIETDMVSEIDPSSSDSGVISRCAVVKPLKFRLCQPIFGKRVGKGRNKNNNARGNCAGVGPAAMGGQAISISAKQNDSACSELEPALPKPKPRDPEKEKRRIARKKEKRATLILGLIMGSFIACWLPFFFLYILVPVCKDCHIPDFAFSLAFWLGYMNSTLNPAIYTIFNKDFRRAFRRILFK*

**Figure S2b.** Nucleotide sequence of the α2 octopamine receptor (AaOAα2-R) open reading frame cloned from *Aedes aegypti* and deduced amino acid sequence. The primer used in RT-qPCR were highlighted in red.

**Figure S2c.**

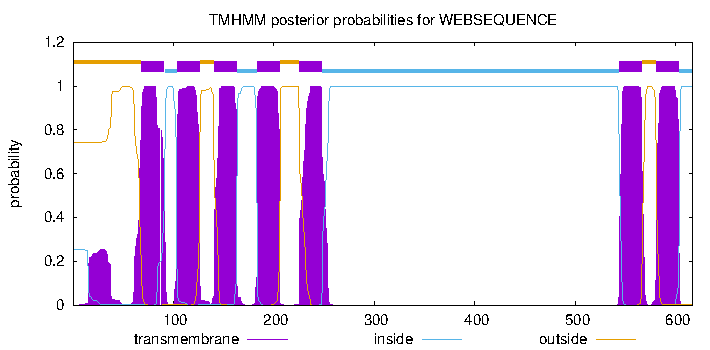

**Figure S2c.** Prediction of the AaOAα2-R transmembrane segments obtained with TMHMM v. 2.0 software.

**Figure S2d.**

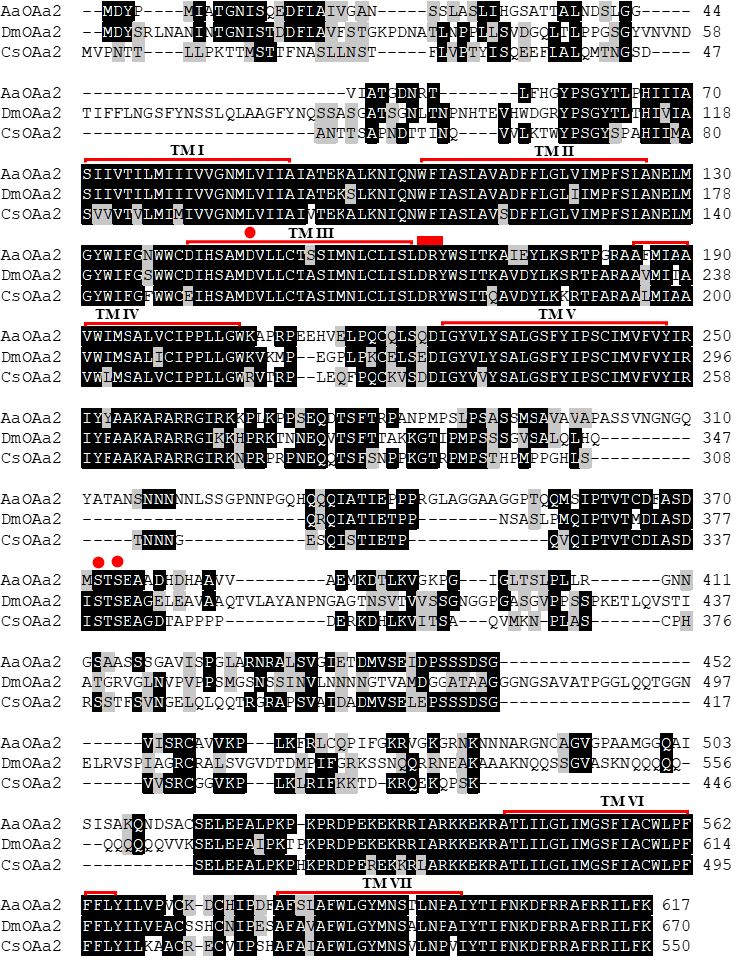

**Figure S2d.** Amino acid sequence alignment of AaOAα2-R with orthologous receptors from *D. melanogaster* (DmOAα2-R) and *C. suppressalis* (CsOAα2-R). The putative seven transmembrane domains (TM I–VII) are indicated by red lines. Identical residues are highlighted in black while conservative substitutions are in grey. A red dot indicates the conserved aspartic acid and the serine residues that could interact with TA.
