## Supplementary material for "Octopamine and tyramine signaling in *Aedes aegypti:* characterization, distribution and potential role in the Dengue vector development and physiology": S3. AaOAb2-R.docx

**Supplementary figures 3.** *Aedes aegypti* β2 octopamine receptor (AaOAβ2-R).

**Figure S3a.**

CGACGGTTTTTCGACATTTCGTGGACGTCGTGTGTCTCATCTTCTTTCCCTTGGTGGCTTTGGTTGGCTTGAATGAGGGGTGGTTTTGGCTCCACGACGACGACGACGACGACATCCACAGCGATTGACTGGTGAGCTCCTAGCTCAGCCCGAGATTGGCGACGCTCTTTGGTCCCAGTTCTATTCGGTCAGCACTCGCAGTCTGTCGCGCAAAACTTCTCTACTAGGGCTCGGTGGACCTTCCGTGTGGTGTGACGGTGTCGCGTGCTATTCAGCTTCCCAGCAAGCAAACGCGGGGACTCCTTGCAGAACGTTCAGGGGGTGGAGAGGTGGCCCTGAGTGTGTTTTCGGTGGGTGGTTTAAGAAGGAGGTGGTTGTGGTGCCGCGTCTCGCGTGTTTTGGGGGGGCTTTTCATATGCTCATTCAAGGGAACGCATAGTGTTGTGCCAACATACAGATCTTGACGAAAATTGGGTGTTTGCGGGAAGGGATAAAGGAACTTCCGGTTGTTCTTATTTTTTCGGGATAGTGTTTCGCAGAAGAGTGTTAGGGCAATTGCGAAGACGAAGCGTTTGGGGTTGCTGTCATCGATTTACGGCACTACGAAGAAATTAGCTGTATTTTGTGTGGTGATTGATAAAAGCAGAGGTTGCAAGGTGATTGGAGAGAAGAGGCTCCACTAGGAGGAGCGAAGAAGCATGCTGTGAACTAAAACAGACATGGAATCACTTGTGCGGCGGTTTTACGGTTTTCCACAGACAGTGCTTGAAATGTAATTAATTTCGCAAATTTCGTTCGTGTTCCAGCCCGGTACCAGCTGGGTGCTCACTTGGTAATCGCTGAATCCATCTAGCAATAAATGCCGAATTCCCCGTTTGAATAGCCTACATCAAATTAGCAAACGCAAAGCAAATTCGAAAAGTGGTGGTTGTGAAAAGTTATAAGATTAGTTAACTACGTCCACGTGTGTGTGTGTGCCCGTGTGTTGACCTCGTCGATGTTCGTTTGTTAATTAGAGTCCCAATTAGTGGTCGTTGGAGGTGTGCAAGCGAACAAGAGTCGGATCAGTTCGCCCGTTGT**ATG**ATGAATCCTTCCAATGACGACCTACCGAAATACATGTACTCAGCGGCAGCGGCAGCAGTTTCCTCTTCCTTCTCGTTGGCCTCCAGTTCGGGAAGTGCACACAGCCAAGAGGGTCATTCTTCCTTCGGAGGAGGCAGCAACTCCACCGGTGGTACTTCGCTGTTCTTCAATGGAAGTGGCGCGGCACTGGCTTCGGCGGCGGCTAAAGCGGCCGGTTCGTCGACGATTTTCACAGTCGGAGCTAGTGCAGTTCCTGGCCGTGCGGGTGATCCTGGAGGACCGGGTGATCCAACGGGGAATCACGTAATCTTCCAGGGCTACAGCACCAGTTCACCGACTTCGTTGGATATGACGACCGTCGGAAGCGATGCAGTTATGGACGATTGCTGTGGTGCGGCTGCTGCCAGTGGCGAGTGGCTGGACGTGCTGCTGCTGGTGCTGAAGGCATCGATCATGATGTTCATCATCGTGGCGGCCATTTTCGGCAACCTGCTCGTCATCATATCGGTTAAGCGACATAGGAAACTAAGAGTTATAACAAATTATTTTGTGGTATCGTTGGCCATGGCAGACATCATGGTTGCGATGATGGCCATGACATTTAATTTCAGTGTACAAATTACGGGAAGTTGGAAATTTGGATCCTTCATGTGTGACGTGTGGAACAGTTTGGATGTGTACTTCTCGACCGCCAGTATACTGCATCTGTGCTGTATTTCGGTGGATAGGTATTATGCGATAGTGAAACCACTTAAGTATCCCATAAGCATGACTAAGCGAGTCGTGGCCATAATGTTACTCAATACCTGGATATCACCAGCACTGTTGTCATTCGTACCGATCTTCGCCGGTTGGTACACCACGGACAAGCACAAGGAGGACGTGCTGAACAACCCGGACAGCTGTTTGTTCATCGTGAACAAACCGTACGCCGTCATCTCCAGTTCGATCTCGTTCTTCATCCCCTGCACGATCATGGTGTTCACCTACTTCCAGATATTCCGCGAGGCCAACCGACAGGAGAAGCAGTTGGCCATGCGCCAAGGGACTGCCATGCTGATGCACCAGCACAACGCCGGTGGCGGAGGCGTAGGAGGAACCGGATCGGGAAGTGCCGGAGGAGGGCGTACGAACGGCGAAGCCCTCAGCGGATCCGGCTCGTCCCGGACGCTCACCATGCACGAGGTGGATGCCGAACAGACGCCCACCAAGGATCGACATCTGATCAAAATGAAACGAGAACATAAGGCTGCACGAACGCTCGGAATCATCATGGGAACGTTTATCCTCTGTTGGCTTCCGTTTTTTCTATGGTACATAATCACGTCGCTCTGCGACGAATGTCCCAATCCGGACATCGTGGTGGTCCTAGTTTTCTGGGTCGGCTACTTCAACTCGACGTTGAACCCGCTGATCTACGCCTACTTCAACCGGGACTTCCGGGAAGCGTTCCGAAACACGCTGGACTGCATGTTCTGCGCGTGGTGGCGCCGGGAAACGTCCCCGCTGGACATCAACGTGCGGAGGTCCAGCTTGCGATACGACTGCCGAGCGAGGAGCGTCTACTCGGAGAACTACCTCCGATCGACGACGCAGAATGACCGGCTCAACAGCGAGATCGGCGAGAGTCTC**TAA**TGTCTTCTGCCTGCATCGTCCGGGACGTCTCGGGACACCATCGGCACTACGGTTATCATGCATCAGAGTCCCCATCTGTCGTCATCGAAGCAGGAGGAGTGCAGTGCGGTCTAGAGAGGGATGGGTGTGTGGATTGGTGTGATCGGAGACGTCTTCGGTTCGGTGAGGGCCAAGGCTGAACAAAATACCAAAAAGTAGATCAATACACGAAGCTAGGCTAAGCTATTTATTTATTTTTTTAATAGTGTAGTTGTGATAGAACTGTGCGGGTGACAGTGTTGCCACTTATGTTAGGTGCGTATCCGTTGTGACTAATTCAAATTGGATAATAATAGGTGAACTGGATGGAAACACTGATAGAGAAGTGTGAGCGAATGCGCAGGTAGAGTGCTAGGAAGAGTTTCAAACACTTGGTAGTGTGATAGCCAACGTTAGTATCGTAGTAGATAACTATACTTGTAGATTAAATGAGACATTCAATTATAATGCCAGGAAAAATGTACTTCGATTCGAAGAAATGGGAGAAAATGACGTTTGTTCGGTCTACGAGGATAGAGCATCGCGAAAGGAAGACAGATTTCATCAATGCAGTGGTTA

**Figure S3a.** *Aedes aegypti* β2 octopamine receptor (OAβ2-R) complete annotated sequence (XM_021837650.1). The primer used to amplify OAβ2-R were highlighted in blue. The start and the stop codon were highlighted in yellow.

**Figure S3b.**

ATG ATG AAT CCT TCC AAT GAC GAC CTA CCG AAA TAC ATG TAC TCA GCG GCA GCG GCA GCA GTT TCC TCT TCC TTC TCG TTG GCC TCC AGT TCG GGA AGT GCA CAC AGC CAA GAG GGT CAT TCT TCC TTC GGA GGA GGC AGC AAC TCC ACC GGT GGT ACT TCG CTG TTC TTC AAT GGA AGT GGC GCG GCA CTG GCT TCG GCG GCG GCT AAA GCG GCC GGT TCG TCG ACG ATT TTC ACA GTC GGA GCT AGT GCA GTA CCT GGC CGT GCG GGT GAT CCT GGA GGA CCG GGT GAT CCA ACG GGG AAT CAC GTA ATC TTC CAG GGC TAC AGC ACC AGT TCA CCG ACT TCG TTG GAT ATG ACG ACC GTC GGA AGC GAT GCA GTT ATG GAC GAT TGC TGT GGT GCG GCT GCT GCC AGT GGC GAG TGG CTG GAC GTG CTG CTG CTG GTG CTG AAG GCA TCG ATC ATG ATG TTC ATC ATC GTG GCG GCC ATT TTC GGC AAC CTG CTC GTC ATC ATA TCG GTT AAG CGA CAT AGG AAA CTA AGA GTT ATA ACA AAT TAT TTT GTG GTA TCG TTG GCC ATG GCA GAC ATC ATG GTT GCG ATG ATG GCC ATG ACA TTT AAT TTC AGT GTA CAA ATT ACG GGA AGT TGG AAA TTT GGA TCC TTT ATG TGT GAC GTG TGG AAC AGT TTG GAT GTG TAC TTC TCG ACC GCC AGT ATA CTG CAT CTG TGC TGT ATT TCG GTG GAT AGG TAT TAT GCG ATA GTG AAA CCC CTT AAA TAT CCC ATA AGC ATG ACT AAG CGA GTC GTG GCC ATA ATG TTA CTC AAT ACC TGG ATA TCA CCA GCA CTG TTG TCA TTC GTA CCG ATC TTC GCC GGT TGG TAC ACC ACG GAC AAG CAC AAG GAG GAC GTG CTG AAC AAC CCG GAC AGC TGT TTG TTC ATC GTG AAC AAA CCG TAT GCC GTC ATC TCC AGT TCG ATC TCG TTC TTC ATC CCC TGC ACG ATC ATG GTG TTT ACC TAC TTC CAG ATA TTC CGC GAG GCC AAC CGA CAG GAG AAG CAG TTG GCT ATG CGC CAA GGG ACT GCC ATG CTG ATG CAC CAG CAC AAC GCA GGT GGC GGA GGC GTA GGA GGA ACC GGA TCG GGA AGT GCC GGA GGA GGG CGT AAC AAC GGC GAA GCC CTC AGC GGA TCC GGC TCG TCC CGG ACG CTC ACC ATG CAC GAG GTG GAT GCC GAA CAG ACG CCC ACC AAG GAT CGA CAT CTG ATC AAA ATG AAA CGA GAA CAT AAG GCT GCA CGA ACG CTC GGA ATC ATC ATG GGA ACG TTT ATA CTC TGT TGG CTT CCG TTT TTT CTA TGG TAC ATA ATC ACG TCG CTC TGC GAC GAA TGT CCC AAT CCG GAC ATC GTG GTG GTC CTA GTT TTC TGG GTC GGC TAC TTC AAC TCG ACG TTG AAC CCG CTG ATC TAC GCC TAC TTC AAC CGG GAC TTC CGG GAA GCG TTC CGA AAC ACG CTG GAC TGC ATG TTC TGT GCG TGG TGG CGC CGG GAA ACG TCC CCG CTG GAC ATC AAC GTG CGG AGA TCC AGC TTG CGA TAC GAC TGC CGA GCG AGG AGC GTC TAC TCG GAG AAC TAC CTC CGA TCG ACG ACG CAG AAT GAC CGG CTC AAC AGC GAG ATC GGC GAG AGT CTC TAA

MMNPSNDDLPKYMYSAAAAAVSSSFSLASSSGSAHSQEGHSSFGGGSNSTGGTSLFFNGSGAALASAAAKAAGSSTIFTVGASAVPGRAGDPGGPGDPTGNHVIFQGYSTSSPTSLDMTTVGSDAVMDDCCGAAAASGEWLDVLLLVLKASIMMFIIVAAIFGNLLVIISVKRHRKLRVITNYFVVSLAMADIMVAMMAMTFNFSVQITGSWKFGSFMCDVWNSLDVYFSTASILHLCCISVDRYYAIVKPLKYPISMTKRVVAIMLLNTWISPALLSFVPIFAGWYTTDKHKEDVLNNPDSCLFIVNKPYAVISSSISFFIPCTIMVFTYFQIFREANRQEKQLAMRQGTAMLMHQHNAGGGGVGGTGSGSAGGGRNNGEALSGSGSSRTLTMHEVDAEQTPTKDRHLIKMKREHKAARTLGIIMGTFILCWLPFFLWYIITSLCDECPNPDIVVVLVFWVGYFNSTLNPLIYAYFNRDFREAFRNTLDCMFCAWWRRETSPLDINVRRSSLRYDCRARSVYSENYLRSTTQNDRLNSEIGESL*

**Figure S3b.** Nucleotide sequence of the β2 octopamine receptor (OAβ2-R) open reading frame cloned from *Aedes aegypti* and deduced amino acid sequence. The primer used in RT-qPCR were highlighted in red.

**Figure S3c.**

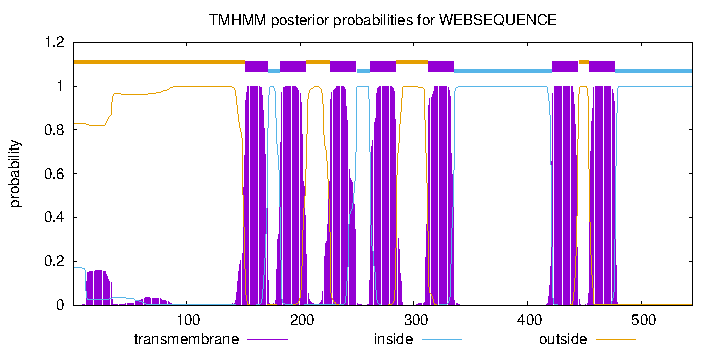

**Figure S3c.** Prediction of the OAβ2-R transmembrane segments obtained with TMHMM v. 2.0 software.

**Figure S3d.**

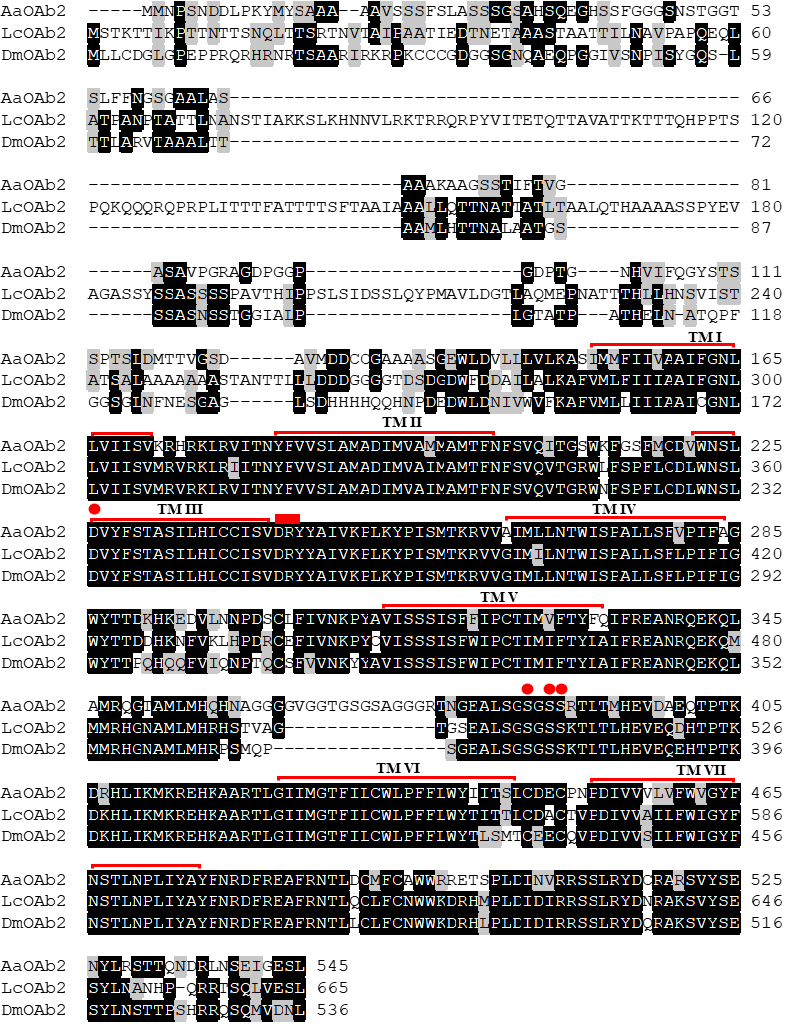

**Figure S3d.** Amino acid sequence alignment of OAβ2-R with orthologous receptors from *L. cuprina* (LcOAβ2-R) and *D. melanogaster* (DmOAβ2-R). The putative seven transmembrane domains (TM I–VII) are indicated by red lines. Identical residues are highlighted in black while conservative substitutions are in grey. A red dot indicates the conserved aspartic acid and the serine residues that could interact with TA.
