## Supplementary material for "Octopamine and tyramine signaling in *Aedes aegypti:* characterization, distribution and potential role in the Dengue vector development and physiology": S4. AaOAb3-R.docx

**Supplementary figures 4.** *Aedes aegypti* β3 octopamine receptor (Oaβ3-R).

**Figure S4a.** CGGTTCGCGAAGCCAGCAAAATTTCAAATCGCGCGCGGTGAATATTCGTGGGACTTCCTAAAATAGTCATTGTGTCTGTGCCATTGCGTGTGCCTAAAGCATCATTATTCAATTTCTGATCTGCATTCGTGAGCACACGAAGTGTCGTTCGGATGCTTCACGCTGATTTTCGTCGCCGTGAAACGGTGGGGGGTTTTGCTCGTCGACCGTTGGGAAAGTGGGGATGCATGCGAAGCAATCTGAGTTGTGGAGAAACCACATGCGGAGAGGGGTGGGGGAGCAAGCACGTTTTGTGACATCCGCATCGGTCGTTATCTGTGAATGGTAGTGAAGTTGCTTTATGGCTACACAGCTAGTCGGAAAGATTGCGGTGATGGTTTTTGGTTGTTTGTCTGAACGGATGGACGCCGACGCCGGTCGGTCAGGAATGTCGACGGTAACAGCAGGGCAACCCGGCGTGTGCTTCGTAGCAAATATGTGATGTGTTGTTTGAACTGGGCGGGAGCCAGTCGGTAGTGGAAGCACCCTGCAGAGTTGCGATAAGTGTCTTGGGAAAAAAGCAAAACCGGAGGCACTCGCAAGTGAAAAAGTAATAATAATTTAATTACCGTCATAATTAGAGCTAAGAGTGTGTGTAATAAATACCGGGAAGGGAAAGTGAGAGTCCGGAAAAAAGGCGAGCCATACGTCGAAGCTCCGAGTTTTGCTCGAGCGTGTGTGTGTGGATGTGTTCTTTATTTTCGTGTTCTTCGAAGAAAGTGTACGGATAAATCGCGTGTGTCAAGGTGAGCAAGTGAAGTTCAATGGCTACGAGGGTCACAAAGTCGGTCGGAGCGTGTTG**ATG**GCCCTCAGAGCAATGTTGGCCGGAGCGGAGATGGTCCCCAAGCCGGTCACGGCAGCTACCCTTGATCCACTGGCAATGGTGACGTCGAACTCGTCCACCCTCGCCGTCGGGAACGTGACTTCGCCGTACCAGTTCTACGGCGGCGGCGGCGGGAACGGAACGCTGTCGGCCCCCGTTACGCCCCTCGTAACGGATCTGATCACGACGGCGGCGGCCAACGGAACGACCACTGCGCTTTCCACGCCCGACGAGGACCGCTTCTTCGAGGTGGGCGTTCTGTGTATGAAGAGTTTAATCTTCGGATCCATCATCATCGGGGCTGTGCTAGGCAATGCACTCGTTATCATATCGGTGCACAAAAATAGAAAGCTAAGAGTAATAACCAATTATTTCGTAGTATCATTAGCAATGGCAGATATCATGGTGGCCCTTTGCGCCATGACCTTCAATGCATCCGTCGAGCTGTCCGGGAGGTGGCTGTTTGGACCGTTTATGTGCGACGTGTGGAACAGCCTGGATGTGTACTTCTCTACGGCCAGCATCCTGCATCTGTGCTGCATTTCTGTCGATCGCTACTTCGCCATCGTGCGACCCCTCGAGTACCCGCTGTACATGACGCACCGCACCGTCGGGTTCATGCTGGCCAACGTGTGGATGCTGCCGGCGTTGATTTCGTTCACGCCCATCTTCCTCGGCTGGTACACGACAGCCGAGCACTTGGAAACGCTCAAGGACAACCCGGAGATGTGCGTTTTCGTGGTCAACAAAGCGTACGCGCTCATCTCCAGCTCCATCAGCTTCTGGATACCGGGAATCGTGATGGTCACCATGTACTATCGGATATTCAAGGAAGCCGTCCGGCAGCGGAAGGCTCTCAGCCGGACCAGTTCGAACATCCTGCTGAACAGTGTTGCCATCAACAACTCCAACACGCTGCACGCCAACTATCACCGCAGTCACCATCTGCGGGCCAGCGATTGCGATCTGGGGATGGACTCAAAGGACATTCAGCACGACACGCACAGCGAAATGAGCGATCTGGATATCCCCATCGTGCCGACTATCATCAACACCGACGGCAACATCGTGGAGAACGACGATGACTTCCTGGTTCCATCGTCGCCACCCCGAAGGCTCAGCCGCAGCAGTATAGACCTGCGAGACTTAGAATACAAGAACGAAACCAAGATCAAACCGTCGGACAGCGTGGGTTCGATCTTTCCGCTGCAGTACGAGAACAACCTGCGCTACTCGGGCAGCGAGCCAATTCACGTCAAAGAAAACCAACAGCGGAGGCCTTCGCGGAAGGAAAGCTTCAAGCAGATCAACTTCTTCCAGCCGTTCTTCTCGAAGAACAGTCTGCTCAAGTCGTACATCAAGGGCGGCAGCTGCAAGGGCGATAACGAAGCAAGCAATCTCAACAATAGCAATAACCGGACGGCTTCCGAGAAGAAGCTGAACAACAAGAACAATGACAATCGGTCCAGCAACAACATAATCGATCTGATCAACGTGACGTCGTCGCTAAACAAGGACAGCTCCGCGGCGGAGAAGGAAGTTAACACGCACAAGCGGCACCAGATTGCGGCCAGCGAGTCGGATTTTCTGACCATGATTCAGAAGCAAAGGTGCCAACGGCTGGGCTTGGGGAAGAACAATGCCAGCTTGAAGAAAAGTAGCAATAAGGATAGATCCAAAGATGATGAACTGATCTATGCGTTAAGCGATTCGGACTTCACGGCTGCATTCCAAGGAGATGAATCAATTTTGAAGAGTTTAAAAAATTTAAAGGCATGTGCCGAGAAGAAAGACGATATTTTTAAGATAGTGTTGAATGGTGATAGTGGAACGGCAGCTACAACGGCAACGTCCGGCTTGAACGATACCCCGGATATCCTAATCGGTTGTTTTGACCACGGGTTCAGCGGAGCCAGACCACTGAGTCCTACCAGTCCCAAAGAGGTTGTGTCCGTTTGTGCAGAGGAAAGCCTCCTCATCAAATCTTTCAATCTCAAAGTTGTGAATAGTAGTAGTAATATGGCTAACCAAAGTAGCACAGTTAGTGATAGTAATCTAAGTCACGCCGAAGGTAGCTTCGTGATGGACCTGATATCTGCAATCGCATCGGCGGGATCTCCACCGCACTCTGCGGGACTGCTTCCGGAAGCAATCTCGCGGGAAAACGATCGCCGCCGAAATTGTGATAAACCACCACCACCTAGCGGTAACAACGTGGACAGTCTGAAACCGTTGGCGACATTGACCGATTCAGGCAGTCCGGTAAACGTCCTGAGCGAAGCATCCACCATATCTCTAGACCTAGTGGCGATGAGTCCGAGCACAAAACTACAAGTTCCTAGCTCTACCGGGGGAACCGGTCTGCAGCCGCCATCCGGTGGTCCAACAAACGAGGACAGTGTCAACCCGGTGGACCATCCGGTCGATCCGGGACAAGTCGTAAACAATCCCGACATCATAATCGAAAATATCCGAAACGACAGCTTAAGCAATCAGCAGCAGCAGACATCGGTGACAGGGACGATTACGATGACGGTTACGACGACGACGACGACAGCAGCGACTCCGATGATGCCACCGTCGATTATGACTCCGTCGCCGATTCACCACCCGGGTCTGCAGCGCAAACGGCAATCGTCCACCGTCACGTACAACATTAACGTGATTAACTTTAGCGATAATCCCGATGATGACAGCAGCTATCAGATGAACAATCGCAGCAGCGAACAGCGGATGGGCGGTGGTGCCGGGATGGGAACCGGTGCCGGTGCGAGATCCAACAGCTCAACGACTTCGAGCCTAAAGCAGCACAACCGCCGCGGCGCCATCTGCATCGTGATCAACAACGAGAATGTGGTTCACGGCACCGACACGGACACGGACTTTGACGAGCACGATGACGCCATGGGCCAGGGTGGCCGTGGTTCCGGAAGTGGTCCACGTCGTTGTGATTGCGGAGGATCCCAGGGCGCGCATCACGACGATGGGTCCGGCGCGGCCAGCTGTCGCCACAACAAGTTCCGTAAGATCAAATGCCTGTGGAACCATTTCAAGCGCCACGGCTTCCTAAGGAGGGGAGGTCTCCACCCGAGGAGGCGCCGAAACGAACCTTGCGACGCTGTTGCGAAGGGCAGACAGAGCTCCGATGGTGGCTCCTCGATGGCTAGGCTCACGGTGCCGACGGGAATCGGTGGCCGCTTCGGTAGTCGTTGTAGCAGTGTTGGTTGTGGGGAAGGTGGCACTTCGACGCGGCAATCGAAGGGATGGCGAGCGGAGCACAAGGCGGCCCGAACCCTGGGAATCATCATGGGAGTGTTCCTGCTCTGCTGGTTACCGTTTTTCCTCTGGTACGTAATCACGACCCTGTGCGGTGAGGAGGCGTGCCCCTGCCCGGACGTGGTCATCACGGTACTGTTCTGGATTGGTTACTTCAACTCGACGCTGAACCCGCTTATCTACGCTTACTTCAATCGGGACTTCCGGGAAGCGTTCCGGAACACGCTCCAATCGCTACTGCCCTGCTTCGGCAAGAAGGACCCCTTCGACGGACACAGCGCCTACTACGTG**TAG**TATTAAGGGTACCCCGGTCCCCGGTCCTGTTCCCGTTTCCCACCTTAGGCTAAGGAGCAAGTTATTGGTGCCGATGGGAGGAATTTAACCCGGTTGGTAGAGATGATGGATCAATCCGTCGTTGGAAAATAAATCGCTCTCTAATCTTT

**Figure S4a.** *Aedes aegypti* β3 octopamine receptor (Oaβ3-R) complete annotated sequence (XM_021837643.1). The primer used to amplify OAβ2-R were highlighted in blue. The start and the stop codon were highlighted in yellow.

**Figure S4b.**

ATG GCC CTC AGA GCA ATG TTG GCC GGA GCG GAG ATG GTC CCC AAG CCG GTC ACG GCA GCT ACC CTT GAC CCA CTG GCA ATG GTG ACG TCG AAC TCG TCC ACC CTC GCC GTC GGG AAC GTG ACT TCG CCG TAC CAG TTC TAC GGC GGC GGC GGC GGG AAC GGA ACG CTG TCG GCC CCC GTT ACG CCG CTC GTA ACG GAT CTG ATC ACG ACG GCG GCG GCC AAC GGA ACG ACC ACT GCG CTT TCC ACG CCC GAC GAG GAC CGC TTC TTC GAG GTG GGC GTT CTG TGT ATG AAG AGT TTA ATC TTC GGA TCC ATC ATC ATC GGG GCT GTG CTA GGC AAT GCA CTC GTT ATC ATA TCG GTG CAC AAA AAT AGA AAG CTA AGA GTA ATA ACC AAT TAT TTC GTA GTA TCA TTA GCA ATG GCA GAC ATC ATG GTG GCC CTT TGC GCC ATG ACC TTC AAT GCA TCC GTC GAG CTG TCC GGG AGG TGG CTG TTT GGA CCG TTT ATG TGC GAC GTG TGG AAC AGC CTG GAT GTG TAC TTC TCT ACG GCC AGC ATC CTG CAT CTG TGC TGC ATT TCT GTC GAT CGC TAC TTC GCC ATC GTG CGA CCC CTC GAG TAC CCG CTG TAC ATG ACG CAC CGC ACC GTC GGG TTC ATG CTG GCC AAT GTC TGG ATG CTA CCG GCG TTG ATT TCG TTC ACG CCC ATC TTC CTC GGC TGG TAC ACG ACA GCC GAG CAC TTG GAA ACG CTC AAG GAC AAC CCG GAG ATG TGC GTT TTC GTG GTC AAC AAA GCG TAC GCG CTC ATC TCC AGC TCC ATC AGC TTT TGG ATA CCG GGA ATC GTG ATG GTC ACC ATG TAC TAT CGG ATA TTC AAG GAA GCC GTC CGG CAG CGG AAG GCT CTC AGC CGA ACC AGT TCG AAC ATC CTG CTG AAC AGT GTT GCC ATC AAC AAC TCC AAC ACG CTG CAC GCC AAC TAT CAT CGC AGT CAC CAT CTG CGG GCC AGC GAT TGC GAT CTG GGG ATG GAC TCA AAG GAC ATT CAG CAC GAC ACG CAC AGC GAA ATG AGC GAT CTG GAT ATC CCC ATC GTG CCG ACT ATC ATC AAC ACC GAC GGC AAC ATC GTG GAG AAC GAC GAT GAC TTC CTG GTT CCA TCG TCG CCA CCC CGA AGG CTC AGC CGC AGC AGT ATA GAC CTG CGA GAC TTA GAA TAC AAG AAC GAA ACC AAA ATC AAA CCG TCG GAC AGC GTG GGT TCG ATC TTC CCG CTG CAG TAC GAG AAC AAC CTG CGC TAC TCG GGC AGC GAG CCA ATT CAC GTC AAA GAA AAC CAA CAG CGG AGG CCT TCG CGG AAG GAA AGC TTC AAG CAG ATC AAC TTC TTC CAG CCG TTC TTC TCG AAG AAC AGT CTG CTC AAG TCA TAC ATC AAG GGC GGC AGC TGC AAG GGC GAT AAC GAA GCA AGC AAT CTC AAC AAT AGC AAT AAC CGG ACG GCT TCC GAG AAG AAG CTG AAC AAC AAG AAC AAT GAC AAT CGG TCC AGC AAC AAC ATA ATC GAT CTG ATC AAC GTT ACG TCG TCG CTA AAC AAG GAC AGC TCC GCG GCG GAA AAG GAA GTT AAC ACG CAC AAG CGG CAC CAG ATT GCG GCC AGC GAG TCG GAT TTT CTG ACC ATG ATT CAG AAG CAA AGG TGC CAA CGG CTG GGC TTG GGG AAG AAC AAT GGC ATC TTG AAG AAA AGT AGC AAT AAG GAT AGA TCC AAA GAT GAT GAA CTG ATC TAT GCG TTA AGC GAT TCG GAC TTC ACG GCT GCA TTC CAA GGA GAT GAA TCA ATT TTG AAG AGT TTA AAA AAT TTA AAG GCA TGT GCC GAG AAG AAA GAC GAT ATT TTT AAG ATA GTG TTG AAT GGT GAT AGT GGA ACG GCA GCT ACA ACG GCA ACG TCC GGC TTG AAC GAT ACC CCG GAT ATC CTA ATC GGT TGT TTT GAC CAC GGG TTC AGT GGA GCC AGA CCA CTG AGT CCT ACC AGT CCA AAA GAG GTT GTG TCC GTT TGT GCA GAA GAA AGC CTC CTC ATC AAA TCT TTC AAT CTC AAA GTT GTG AAT AGT AGT AGT AAT ATG GCT AAC CAA AGT AGC ACA GTT AGT GAT AGT AAT CTA AGT CAC GCC GAA GGT AGC TTC GTG ATG GAC CTG ATA TCT GCA ATC GCA TCG GCG GGA TCT CCA CCA CAC TCT GCG GGA CTG CTA CCG GAA GCA ATC TCG CGT GAA AAC GAT CGC CGC CGA AAT TGT GAT AAA CCA CCA CCA CCT AGC GGT AAC AAC GTG GAC AGT CTG AAA CCG TTG GCG ACA TTG ACT GAT TCA GGC AGT CCG GTA AAC GTC CTG AGC GAA GCA TCC ACC ATA TCT CTT GAC CTA GTG GCG ATG AGT CCG AGC ACA AAA CTA CAA GTT CCT AGC TCT ACC GGG GGA ACC GGT CTG CAG CCG CCA TCC GGT GGT CCC ACG AAC GAG GAC AGT GTC AAC CCG GTG GAC CAT CCG GTC GAT CCG GGA CAA GTC GTA AAC AAT CCC GAC ATC ATA ATC GAA AAT ATC CGA AAC GAC AGC TTA AGA AAT CAG CAG CAG ACA TCG GTG ACA GGG ACG ATT ACG ATG ACG GTT ACG ACG ACG ACG ACG ACA GCA GCG ACT CCG ATG ATG CCA CCG TCG ATT ATG ACT CCG TCG CCG ATT CAC CAC CCG GGT CTG CAG CGC AAA CGG CAA TCG TCC ACC GTC ACG TAC AAC ATT AAC GTG ATT AAC TTT AGC GAT AAT CCC GAT GAT GAC AGC AGC TAT CAG ATG AAC AAT CGC AGC AGC GAA CAG CGG ATG GGT GGT GGT GCC GGG ATG GGA ACC GGT GCC GGT GCG AGA TCC AAC AGC TCA ACG ACT TCG AGC CTA AAG CAG CAC AAC CGC CGC GGC GCC ATC TGC ATC GTG ATC AAC AAC GAG AAT GTG GTT CAC GGC ACC GAC ACG GAT ACG GAC TTT GAC GAG CAC GAT AAC GCC ATG GGC CAG GGT GGC CGT GGT TCC GGA AGT GGT CCA CGT CGT TGT GAT TGC GGA GGA TCC CAG GGC GCG CAT CAC GAC GAT GGG TCC GGC GCG GCC AGC TGT CGC CAC AAC AAG TTC CGT AAG ATC AAA TGC CTG TGG AAC CAT TTC AAG CGC CAC GGC TTC CTA AGG AGG GGA GGT CTC CAC CCG AGG AGG CGC CGA AAC GAA CCT TGC GAC GCT GTT GCG AAG GGC AGA CAG AGC TCC GAT GGT GGC TCC TCG ATG GCT AGG CTC ACG GTG CCG ACG GGA ATC GGT GGC CGC TTC GGT AGT CGT TGT AGC AGT GTA GGT TGT GGG GAA GGT GGC ACT TCG ACG CGG CAA TCG AAG GGA TGG CGA GCG GAG CAC AAG GCG GCC CGA ACC CTG GGA ATC ATC ATG GGA GTG TTC CTG CTC TGC TGG TTA CCG TTT TTC CTC TGG TAC GTA ATC ACG ACC CTG TGC GGT GAG GAG GCG TGC CCC TGC CCG GAC GTG GTC ATC ACG GTA CTG TTC TGG ATT GGT TAC TTC AAC TCG ACG CTG AAC CCG CTT ATC TAC GCT TAC TTC AAT CGG GAC TTC CGG GAA GCG TTC CGG AAC ACG CTC CAA TCG CTA CTG CCC TGC TTC GGC AAA AAG GAC CCC TTC GAC GGA CAC AGC GCC TAC TAC GTG TAG

MALRAMLAGAEMVPKPVTAATLDPLAMVTSNSSTLAVGNVTSPYQFYGGGGGNGTLSAPVTPLVTDLITTAAANGTTTALSTPDEDRFFEVGVLCMKSLIFGSIIIGAVLGNALVIISVHKNRKLRVITNYFVVSLAMADIMVALCAMTFNASVELSGRWLFGPFMCDVWNSLDVYFSTASILHLCCISVDRYFAIVRPLEYPLYMTHRTVGFMLANVWMLPALISFTPIFLGWYTTAEHLETLKDNPEMCVFVVNKAYALISSSISFWIPGIVMVTMYYRIFKEAVRQRKALSRTSSNILLNSVAINNSNTLHANYHRSHHLRASDCDLGMDSKDIQHDTHSEMSDLDIPIVPTIINTDGNIVENDDDFLVPSSPPRRLSRSSIDLRDLEYKNETKIKPSDSVGSIFPLQYENNLRYSGSEPIHVKENQQRRPSRKESFKQINFFQPFFSKNSLLKSYIKGGSCKGDNEASNLNNSNNRTASEKKLNNKNNDNRSSNNIIDLINVTSSLNKDSSAAEKEVNTHKRHQIAASESDFLTMIQKQRCQRLGLGKNNGILKKSSNKDRSKDDELIYALSDSDFTAAFQGDESILKSLKNLKACAEKKDDIFKIVLNGDSGTAATTATSGLNDTPDILIGCFDHGFSGARPLSPTSPKEVVSVCAEESLLIKSFNLKVVNSSSNMANQSSTVSDSNLSHAEGSFVMDLISAIASAGSPPHSAGLLPEAISRENDRRRNCDKPPPPSGNNVDSLKPLATLTDSGSPVNVLSEASTISLDLVAMSPSTKLQVPSSTGGTGLQPPSGGPTNEDSVNPVDHPVDPGQVVNNPDIIIENIRNDSLRNQQQTSVTGTITMTVTTTTTTAATPMMPPSIMTPSPIHHPGLQRKRQSSTVTYNINVINFSDNPDDDSSYQMNNRSSEQRMGGGAGMGTGAGARSNSSTTSSLKQHNRRGAICIVINNENVVHGTDTDTDFDEHDNAMGQGGRGSGSGPRRCDCGGSQGAHHDDGSGAASCRHNKFRKIKCLWNHFKRHGFLRRGGLHPRRRRNEPCDAVAKGRQSSDGGSSMARLTVPTGIGGRFGSRCSSVGCGEGGTSTRQSKGWRAEHKAARTLGIIMGVFLLCWLPFFLWYVITTLCGEEACPCPDVVITVLFWIGYFNSTLNPLIYAYFNRDFREAFRNTLQSLLPCFGKKDPFDGHSAYYV*

**Figure S4b.** Nucleotide sequence of the β3 octopamine receptor (OAβ3-R) open reading frame cloned from *Aedes aegypti* and deduced amino acid sequence. The primer used in RT-qPCR were highlighted in red.

**Figure S4c.**

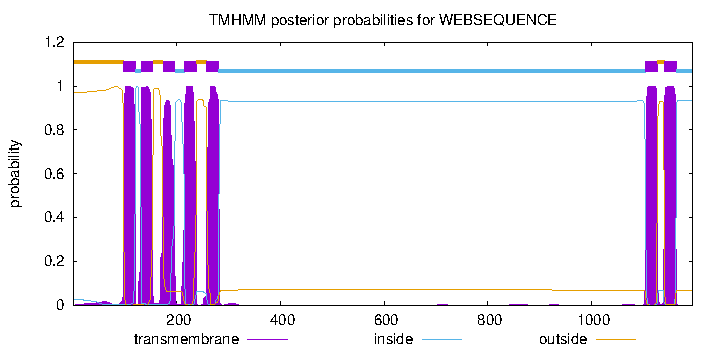

**Figure S4c.** Prediction of the Oaβ3-R transmembrane segments obtained with TMHMM v. 2.0 software.

**Figure S4d.**

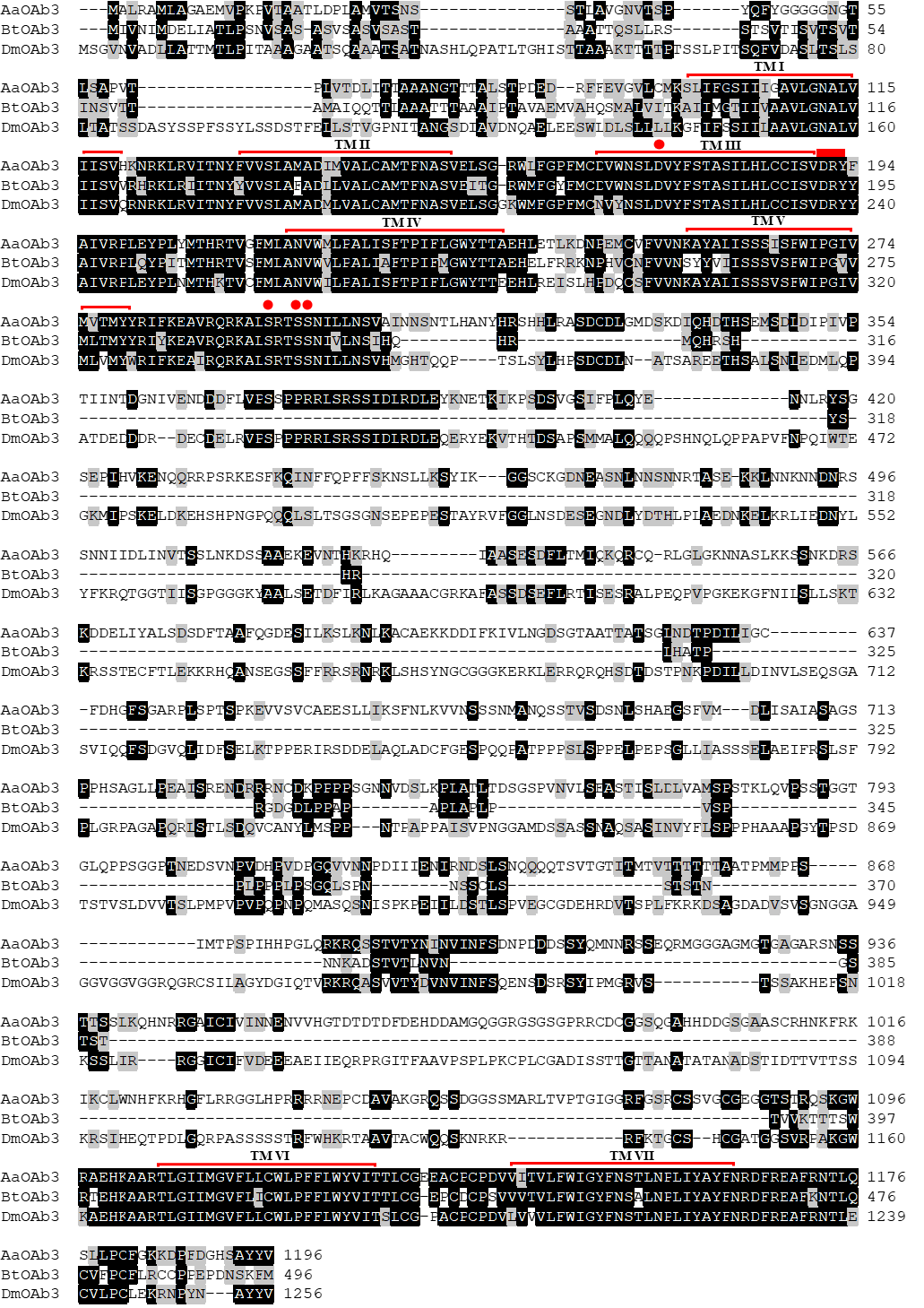

**Figure S4d.** Amino acid sequence alignment of Oaβ3-R with orthologous receptors from *B. tabaci* (BtOAβ3-R) and *D. melanogaster* (DmOAβ3-R). The putative seven transmembrane domains (TM I–VII) are indicated by red lines. Identical residues are highlighted in black while conservative substitutions are in grey. A red dot indicates the conserved aspartic acid and the serine residues that could interact with TA.
