## Supplementary material for "Octopamine and tyramine signaling in *Aedes aegypti:* characterization, distribution and potential role in the Dengue vector development and physiology": S5. AaTAR1.docx

**Supplementary figures 5.** *Aedes aegypti* type 1 tyramine receptor (AaTAR1).

**Figure S5a.**

TGCGATCGTTATCGAAATCTAAAACAATTTACCTAATACCTATAACGTGAACGAATAGAAAGTTCATTTTTCGGATAGCTTGTGCTAGCCTGCACATCTTGATTAGCGTTTATGATATACTCTACCACCAACCAAAACATATTCGAGTTCATTTCATTCATAAAATTGGCACTACGTGAACCCGGTTTTCCCCGCTGCAATGCTTTTCGCGCCCAAAAGAAAACTCTCTCCACTCATTTCCACTCGCTCTCGCGGCAACTCATTTCCCACATGGCCCACTCACTCGTCGTTGATCCCGCATGCATATTCGCGCAGGTTATCGCTCGATGATTGCTCTTTCTTGACGATTGCCACACCATCCGGTCGGAGATACAAACGCCAGAACAGACACTTCCACGGTCCTCAGATGGACCAACCAGTCGTCAGTCAACGTCGGAATGTGGTTGTAAGCGCACCGAAAATCCTCCGGTATCTCCCCATCAGCTCCGAATAGGGTTCTGTATTGGGTCTCTTCTCACTGGATAAAACTGACTGTTGATGATTATTACTGTGTTAAATCTCCAGATGTACTCGAAGTGCGGAAAATGGGATGCCGCCATCCGGTAGTTTAGCAGTGTGATTGAATTTGTTTTCACCAAGAGAAGCGTGTTCAACTGTTCGGGCGGTTGGGGCGTTTCTCCAAACAGGAAAAAGTAAAGTGTTTACGTGGAAAAGCGCGAAGGTTAACACTCAAGTGAGGCGAAGTGCGCTCGATAGCGGCGTTTTTCCCACGGCTTCGATGTTTCTTCAAAGTGTGTTTTGCCATTGACAAGAAATGTGATTTCCGAGAAAAAGGGGGAACAATTTACTATGCCGATGTTGACGAAGATGGACGGAGTCTGGGCAGAAAAATGTGATTGCACCACCATCAGGTCGTATTAGTAGGCTGTGAAAAACGTGAACGCGAGTGATGATAACGATGAAAATTCGCACCTGCTGAGAAATTCGGTGGAGAAAAACGACGGGTGGGAAATGTGTTTACGAAAAAGTGGAAGAAAATACGCGAATGAATTAATTTAAAAGAGTGCATATTTTCCCATTACCAGAACAAACACGTTAGCATTTCATCACATCGTTGGGTAGAATTATGAAGAATCAAATGATGATAATCTACTGTTTCCAAATGGCAAACGGTTTGATCAGTGTTCGCGCTAGCAAATAATTACACAAAACCAGAATATAAAATATCCGGGTAATGTTATTTCTCCAAAGCAAACATCAGTGCCAGCAGTGTCAGGAATAACACGAAAACACGTTCTACAACATCAATAACATGCAACAGTGAATGTGCACGAGGTATAAATAACGCAATTATCGTGTTTTTTGGTCACTTGACCGGCGTGGCGTTTCCCCGAGGAAGTGGTGGTGGCCTGCTGTGATTGAGAATCCGTCACAACCAGAAGGAAAGGGGCAGCAGCCATCCGTCACGGCCAACCGTGATCTCCAAAATTACGCCCGATCGTTCGCGATCAATTACTCTCCGTAGAGTGAACCTCCGTAGAAAATCCCAAGAACTCAACAGGAGTACATCACAGAGAGAGAGAGAGCTATTCGCCATCAACAAACAGCGCCTACCGAGGCCTACTGAGGGTTTTGCCGTTGCTGCCCGTAAATTACCGCCGTTCGACGAAGCCTACTGCGATGAGGACCGGCCACAATGGAAGTACAGCAACAGCAGCTTCGAGAGCAAACGAGCAACGTCTAGTGGTTTACTTTGGTGTTTCGAGACCAGAACAGTGAGAATAATGTGTAGAGA**ATG**GCAATTGTATCAGTGATACCAGTTATCAACATTACAAACTCGAGTGACAATGGTGCGAATGGGACGAATGGAACCGTTGGTGGTGGTGGAGGAGTTGGAGAGTTTGACGATGGGAGTGGGTGCCCGAGGCAAGATGAAATCCTGTATCCGAGTATATTTGGAATTGACCTAGCGGTGCCACAATGGGAGGCCATTGCAACCGCACTCATACTGACGCTCATTATCATCATTACCATCGTAGGCAACGTGCTGGTCATATTGAGCGTATTCACATACAAGCCGCTGAGGATTGTGCAGAACTTCTTCATTGTGTCACTGGCAGTAGCGGACCTGACGGTGGCCATTCTGGTGCTCCCATTCAATGTGGCCTATTCGATACTGGGCCGATGGGAATTTGGCATCCACGTATGCAAAATGTGGCTTACCTCGGATGTCCTGTGCTGTACAGCTTCGATTCTAAATCTGTGTGCAATAGCATTAGATAGGTATTGGGCCATTACCGATCCAATTAATTATGCTCAGAAACGTACCTTGGAGCGAGTGCTGGCATTGATTGCCGGTGTGTGGATACTATCGTTGCTGATCAGTTCTCCACCGTTGATCGGCTGGAACGATTGGCCGGAGCCGGAGAAATTCTCCAGCGAGTTCCCGTGTCAATTAACAAGCAATCAAGGTTATGTCATCTACTCTTCGCTAGGATCATTCTACATCCCCCTGATCATCATGACGATTGTGTACATCGAAATTTACATAGCCACTCGGAGACGTCTCCGGGAAAGGGCCCAAGCATCAAAAATCAACACTTTAGCTAGTAGGTGTATCGGACAGAGTGAAAAAGACACTTGTATGAACCAACCGGACCAGGAATCAATTAGCAGTGAAGCCAACCACAACGAACATCCGCACAATAGCACCAGTAGCAGTAGCAATGAACATCGGTCCCAAAGGAAGCGGAAGAAGAAAGCAAAGGAAAAAAAGGAAGCGAAGGAGGCGGCGAAACGTGCAAAGCAAAATCAACTCCGGATAGCCCTCCGGGATGAAGATTCCGTTACGGAGTGTCCGGAAAATTCGTCCATCAGCGCGAAAGCAAACTGTGATATTAAATCGGCCACCGGTGGTCCCAACGGAATGGCTGCAACCGGTACGACGGCAGAGGGTGAGGATGTCAACGCCTCCGCCACCACCAACGGAGACAAGAATCCGCTACAGCAAAAAGCTACATCTTCGGCGGAAACCCGACTCCGAACAAAACAATCGATCCGAAGGCCCGGCGGCCTTAATCAGTTTATCGAAGAGAAGCAGAAGATTTCGCTTTCCAAGGAAAGGAGGGCGGCCAGGACGCTTGGCATCATCATGGGGGTTTTTGTGGTGTGTTGGTTGCCCTTTTTCCTCATGTACGTAATCCTACCGTTCTGTCCAAGCTGTTGTCCGACGAACAAATTAATTAACTTCATCACTTGGCTGGGATACATCAACTCGGCTTTAAATCCCATTATTTACACGATATTCAATTTGGATTACAGGCGGGCATTCAAACGATTGCTCGGAATCAAACAG**TAG**GCTTTTCGTTTCCATCCATGTTCTCATCTTCTCATGATTCGGTTTGGGGAATGGGATAACTAGCTAGGTATCTATGGCAACCCGGTGTGGTGGCTCAAAGGAACCAACGAATGGCATGTTGCACATTGTGTGGCCTGCTTCTTGGTGGTTGGCAGCTTGAATGTACGCATACAGAGGCACCAATTGAGTAGATAGACATTGTGGGAAACATGCAATCAACGTTGTTATTCGAGATGAACCCGCACTGCATCAGCAATCGAGTCAGATTGCACTAATTCGTGTTTTGTATTATATATCTCTCTTTACGGAATCATTTGATGTTATCTTTTGCACCCGAAGGGCCGATGGAAAATTTCAATAATTGAACCTCCATCAATGGTATCAGATGGGCAAAACTATTTTAACAAAATTAACATTCAACGAGAAATTTCAATTACAGTCAACTTCCCACGTCTCGATTGCAAC

**Figure S5a.** *Aedes aegypti* type 1 tyramine receptor (AaTAR1) complete annotated sequence (XM_001652205.3). The primer used to amplify AaTAR1 were highlighted in blue. The start and the stop codon were highlighted in yellow.

**Figure S5b.**

ATG GCA ATT GTA TCA GTG ATA CCC GTT ATC AAC ATT ACA AAC TCG AGT GAC AAC GGC GCG AAT GGG ACG AAT GGA ACC GTT GGT GGT GGT GGA GGA GTT GGA GAG TTT GAC GAT GGG AGT GGG TGC CCG AGG CAA GAT GAA ATC CTG TAT CCG AGT ATA TTT GGA ATT GAC CTA GCG GTG CCA CAA TGG GAG GCC ATT GCA ACC GCA CTC ATA CTG ACG CTC ATT ATC ATC ATT ACC ATC GTA GGC AAC GTG CTG GTC ATA TTG AGC GTA TTC ACA TAC AAG CCG CTG AGG ATT GTG CAG AAC TTC TTC ATT GTG TCA CTG GCA GTA GCG GAC CTG ACG GTG GCC ATT CTG GTG CTC CCA TTC AAT GTG GCC TAT TCG ATA CTG GGC CGA TGG GAA TTT GGC ATC CAC GTA TGC AAA ATG TGG CTT ACC TCG GAT GTC CTG TGC TGT ACA GCT TCG ATT CTA AAT CTG TGT GCA ATA GCA TTA GAT AGG TAT TGG GCC ATT ACT GAT CCA ATT AAT TAT GCT CAG AAA CGT ACC TTG GAG CGA GTG CTG GCA TTG ATT GCC GGT GTG TGG ATA CTA TCG TTG CTG ATC AGT TCT CCA CCG TTG ATC GGC TGG AAC GAT TGG CCG GAG CCG GAG AAA TTC TCC AGC GAG TTC CCC TGT CAA TTA ACA AGC AAT CAA GGT TAT GTA ATC TAC TCT TCG CTA GGA TCA TTC TAC ATC CCC CTG ATC ATC ATG ACG ATT GTG TAC ATC GAA ATA TAC ATA GCC ACT CGG AGA CGT CTC CGG GAA AGG GCC CAA GCA TCA AAA ATC AAC ACT TTA GCT AGT AGG TGT ATC GGA CAG AGT GAA AAA GAC ACT TGT ATG AAC CAA CCG GAC CAG GAA TCA ATT AGC AGT GAA GCC AAC CAC AAC GAA CAT CCG CAC AAT AGC ACG AGT AGC AGT AGC AAT GAA CAT CGA TCC CAA AGG AAG CGG AAG AAG AAA GCA AAG GAA AAG AAG GAA GCG AAG GAG GCG GCG AAA CGT GCA AAG CAA AAT CAA CTC CGG ATA GCC CTC CGG GAT GAA GAT TCT GTT ACG GAG TGT CCG GAA AAT TCG TCC ATC AGC GCG AAA GCA AAC TGT GAT ATT AAA TCG GCC ACC GGT GGT CCC AAC GGA ATG GCT GCA ACC GGT ACG ACG GCA GAG GGT GAG GAT GTC AAC GCT TCC GCC ACC ACC AAC GGA GAC AAG AAT CCG CTA CAG CAA AAA GCT ACG TCT TCG GCG GAA ACC CGA CTC CGA ACA AAA CAA TCG ATC CGA AGG CCC GGC GGC CTT AAT CAG TTT ATC GAA GAG AAG CAG AAG ATT TCG CTT TCC AAG GAA AGG AGG GCG GCC AGG ACG CTT GGC ATC ATC ATG GGG GTT TTT GTG GTG TGT TGG TTG CCC TTT TTC CTC ATG TAC GTA ATC CTA CCG TTC TGT CCA AGC TGC TGT CCG ACG AAC AAA TTA ATT AAC TTC ATC ACT TGG CTG GGA TAC ATC AAC TCG GCT TTA AAT CCC ATT ATT TAC ACG ATA TTC AAT TTG GAT TAC AGG CGG GCA TTC AAA CGA TTA CTC GGA ATC AAA CAG TAG

MAIVSVIPVINITNSSDNGANGTNGTVGGGGGVGEFDDGSGCPRQDEILYPSIFGIDLAVPQWEAIATALILTLIIIITIVGNVLVILSVFTYKPLRIVQNFFIVSLAVADLTVAILVLPFNVAYSILGRWEFGIHVCKMWLTSDVLCCTASILNLCAIALDRYWAITDPINYAQKRTLERVLALIAGVWILSLLISSPPLIGWNDWPEPEKFSSEFPCQLTSNQGYVIYSSLGSFYIPLIIMTIVYIEIYIATRRRLRERAQASKINTLASRCIGQSEKDTCMNQPDQESISSEANHNEHPHNSTSSSSNEHRSQRKRKKKAKEKKEAKEAAKRAKQNQLRIALRDEDSVTECPENSSISAKANCDIKSATGGPNGMAATGTTAEGEDVNASATTNGDKNPLQQKATSSAETRLRTKQSIRRPGGLNQFIEEKQKISLSKERRAARTLGIIMGVFVVCWLPFFLMYVILPFCPSCCPTNKLINFITWLGYINSALNPIIYTIFNLDYRRAFKRLLGIKQ*

**Figure S5b.** Nucleotide sequence of the type 1 tyramine receptor open reading frame cloned from *Aedes aegypti* and deduced amino acid sequence. The primer used in RT-qPCR were highlighted in red.

**Figure S5c.**

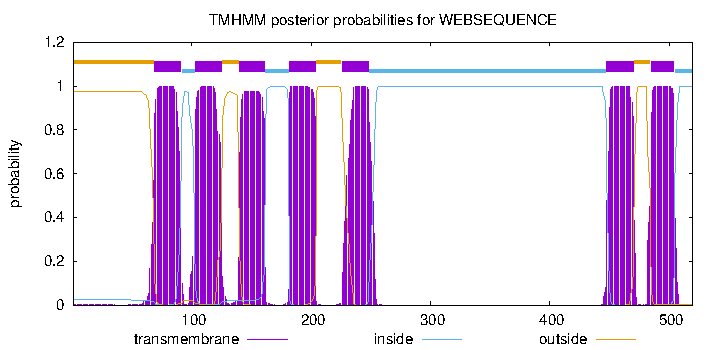

**Figure S5c.** Prediction of the AaTAR1 transmembrane segments obtained with TMHMM v. 2.0 software.

**Figure S5d.**

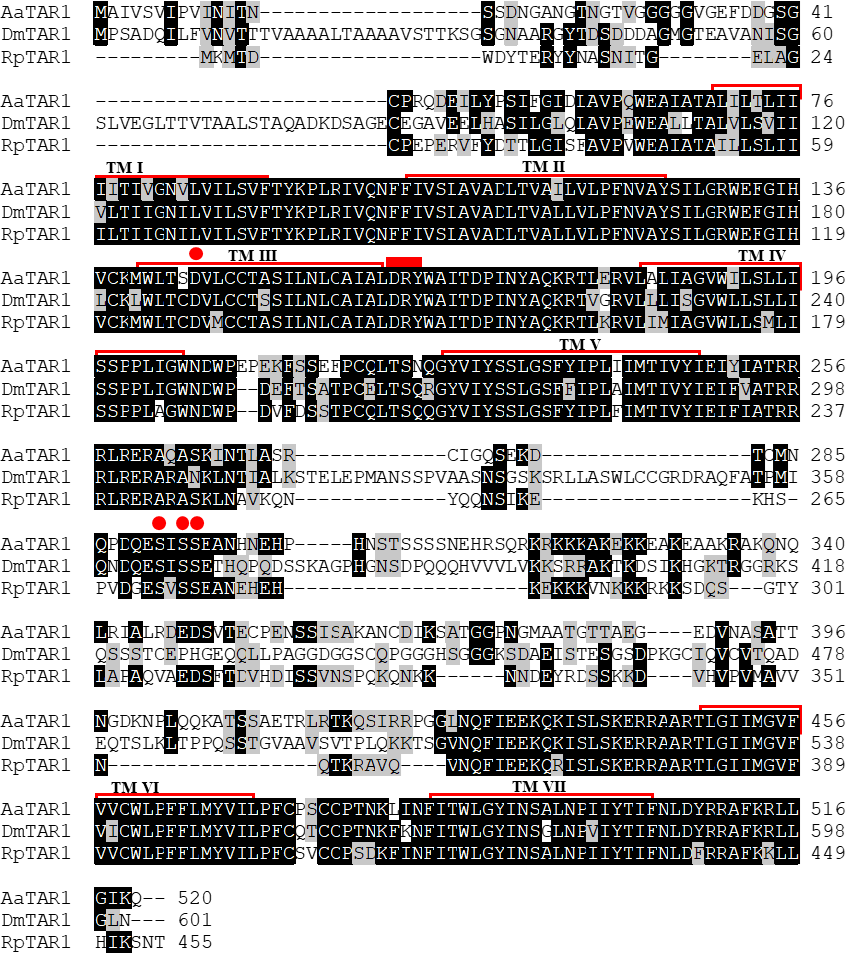

**Figure S5d.** Amino acid sequence alignment of AaTAR1 with orthologous receptors from *D. melanogaster* (DmTAR1) and *R. prolixus* (RpTAR1). The putative seven transmembrane domains (TM I–VII) are indicated by red lines. Identical residues are highlighted in black while conservative substitutions are in grey. A red dot indicates the conserved aspartic acid and the serine residues that could interact with TA.
