## Supplementary material for "Octopamine and tyramine signaling in *Aedes aegypti:* characterization, distribution and potential role in the Dengue vector development and physiology": S6. AaTAR2.docx

**Supplementary figures 6.** *Aedes aegypti* type 2 tyramine receptor (AaTAR2).

**Figure S6a.**

TGATAAAGGGCTCTCGCACAGACCGTTCCCGGAGCTGGAGCGAGAAGCAGCTGGAAACCGAGATTCAACCGCGCGGTAGACTGGTGGGTACTATTGTTGAGTTGCCACCGCACAGTACCCTAAGGTAAACACTACAAAGTGGATGTTCGCCACCGCTTGGTTAGCTGCAGCGGAGTACAGTTCGATTCGAACCTTCATCCGGATTGGTCGCCTGTCGTCCGAAGGACCTCTCGGAAGTGAAACCGCGGCGCGCCGCGTTCTCCATAGGACTATTCTTAGTGATTTTGGGTGTTTAGGTCAGTGTCTGCTGGAGGCCTTCACCTAGAGCGCAGTGCTGTAAATCACTGTTATTTTTTGCTCGCGGAATGTGAAATGGTTTGGAGTGACCCGGTTCCTGTTGTCCCGTTGGAAGCAGTTGGATTAATGCGTTTTTGGTGACCTTTTGGGAGTGAATTTATTACTGAAGCAGTGACGTTTGGTGCTTTCGGCGAGACGAAGCTGACTTTGTGAGGAATCGCGCAGTTTGATAAATAAACAACCAACGGAAAGCGAAGAATTTGTGCGAGTGAGCTCAGAGCTTTAGAGGGGGGCCCGCAATTTGGTGATCTGTGGTGAATGAGCAATTCTAGTATCAACAACAATAACGAAACGAGTGAATAAACAAATAAATATAGCAAGTGAAATATAATACGAGACGCTGTGTCCAGTGTGGTATCTGCCGCCCAGAACGATTGGTTCTAGCTACGAGAGTGCGAAGCGAAACCAATGGGATGGAATAAATGTTGAACGGCGATTGAGACGTACGCGGCCACAGTGGAATGTGACACCTGTTTAAAAGTGATCTACCAAGCGAGGGACTACTATTGAGTTTTTGCGGTACTGTTTGCTCTGATGTGAAACTGGAATAACCTTCAGAAATTGGCGCTGTTTTTCCTAACGATTATTGCGATAATATTCAATGGATTGGTGATCTGCTGTAACGGGAGTATACCCACGCGTGTAGTACTAGAATAGACACAGTATGTATCAGTCGTCGACTACCGAAGCAATGCCGCTGAGTGACCACCAACACCACCACCACTAAG**ATG**GAAACTCGCCTCGATGACCGATGCGCCGCTCTACTGCTTCAACTCAGCGCCGACGACGGAGACTACGCTCAGCAGCTTCCCGAGTATCCTTCCGTCGCAGCGAACTCCAGTACTTCGGAATCACACGATCAACAGTCCCTACTGTTGGTGCCAACGATCGCAACGAAGCTGTCGAAGTTGCTGACCGACTTGGTCAGTGTTGGTGGGGGAACCGGGGGAGTGGCCGGCAGTGTCAATTTCAGTGCATTGTTGTCGCTCAATTTCAGCACCAGTGGCAATAGCAGCTCTATTGGTGTCAGTGCAACAGCAACAACGGCCTTAGCGGTGGCGTTTGACGATCATCTACCATCATCGGAACGCACGCAGAACCACTCCACCGATGGAAGGTTCAACTGCGACGGTCGCAACCTGGCGGAATTGGCAACGAGTGCGTTCGGAAGTGCGACCGGTGGGGGCGCGAGTGGCGATGGGGGGTTCGGTCCGAAATTGGAAGTTTCGAAAACGATGGACGTGATCGGGAACAGTGGTGGTGTTGGTGATGGAACCGATTCGGTCGGGTTCGGCATCGGAGACGGCGGTGCGGCCACGCAACTGTTGATTGCCAACTGGCAGGACTGTGTCGTTGTGATATTATTTTGCTTACTGATCGTAGTGACTGTGATCGGCAACACGCTCGTGATCCTGTCGGTGATTACCACTCGGCGGCTGCGGACCGTCACCAACTGCTTCGTGATGAGCCTGGCCGTGGCCGATTGGTTGGTCGGGATTTTCGTGATGCCCCCGGCTGTCGCGGTCCATCTTTTGGGATCGTGGCCGCTCGGATGGATACTGTGTGATATATGGATCTCATTAGATGTTCTACTGTGTACGGCGTCAATCCTTAGCTTATGTGCAATTAGTATAGATAGATATCTAGCGGTAACGCAACCGCTCAACTACTCTCGCCGACGGCGCTCGAAACGGCTGGCCCTCCTGATGATTCTGGTCGTCTGGGTACTAGCACTGGCCATCACCTGCCCTCCCATTCTCGGCTGGTACGAACCGGGCCGGCGGGAGCTCACCCAGTGCCGGTACAACCAAAACGAGGGCTACGTCGTCTTCTCGGCGATGGGATCCTTCTTCATTCCGATGGCCGTCATGATCTACGTGTACGTGCGGATATCCTGCGTGGTGGCATCCCGCCACGACAAAATGACCGAAATCGAAGTGCACAAGAAAAGTCACCGAACGCGTGATGCCGACGAGCCGTGCTACCACTACGCTTCGGAGATGGATCCGGCCCAGTTGAGTCCCAAGCGAAAGCGTTCCTCATCGCAGCTCTCGACAGCAACCACAACTTACGTGACACAGCTGGGGCCAGGCGGTGGCGGAGGCTCATCCAGCTTTACGGCCCTTAGCGGAGACTGCCCTGGAGGAGATTGCCTTAAGCTGGGGGGAGGCGGCACCAGAAAAAATCACAAGAATGGACATTACGAGCTTGTCGACATCAATTCCATGTCCAAAACCACAACTTCTTTCGTGGGCAACGGTGGAGGTACGAACGGAGCTGGACTCGATGAGGACCATATGGACCCGGGTGGCCAGCGGGAGTCACCGGGAGCTACCGGAGGCATAAAACGCAACTCCACCATCCACTACAAGCTGGCCTCGACGACGGAGCTTTGGTCCTCGGCCAATAACGTTTTCAATCAACAGCAGAAACCACAAACGGCAAATCAATCGGGCGGCTTGAGGCGTACCAATACCATTAGTAACTCAATTACGTCGTCGACCGCCACGGCATCGGCCTCGGCAGTGGCGGCCGGGCGAAATATCCGGATCCACCAGAAGTCACTTTCGTCGCGAATTTTGTCCATGAAGCGCGAGAACAAAACGACCCAAACGCTCAGCATCGTGGTCGGAGGCTTCATCGCGTGCTGGCTGCCCTTCTTCATCCATTACATCATTACGCCGTTCCTGCCAGAAGAGCTGGCTGCACCCAAACTGGGGGAGTTTTTCACATGGCTCGGCTGGATCAACAGCGCCATCAATCCGTTCATCTACGCGTTCTACAGCGTGGACTTCCGGGCGGCCTTCTGGCGCTTGACGTTGCGGCGCTTTTTCCGCAACAGCGAGAAGGCACCCTTTGCCAACATTCACAACATGTCCATGCGGAGA**TAA**GCTCCGACCGCATCCAATTTCCTATAATAGCCGATCCACCCTCTCTCCATCCACAGTTCAAATGACACCTTGTTGCTTGTATAAGCGGAACCTCCGAATCAGAGCCCGGGCAGGTTGAAAAGTGAACGACAAATTTGCAGTTTCTGTCTCGTGTGGCTGGGAAAGTGCGATTTGAGATGTGGTTTCACACGGATGATTCCGCTTCTATTGTCAACACATTATGGGATTACGTACGGATAATATGCTTCGTATGTGAGCGCGATAATGGGCGTTTTGTCTCTGACCTATTGTTATTGGAAAAATGTGATTGTACATTCTGATAAAACGTGTGCTGTACCATTGTGAATCTGAGTAGGTGAATATTCAGTTGGTTATGTTAAAGAACTAAAGAACAGATTTTAATAACTTA

**Figure S6a.** *Aedes aegypti* type 2 tyramine receptor (AaTAR2) complete annotated sequence (XM_021837305.1). The primer used to amplify AaTAR2 were highlighted in blue. The start and the stop codon were highlighted in yellow.

**Figure S6b.**

ATG GAA ACT CGC CTC GAT GAC CGA TGT GCG GCT CTA CTG CTT CAG CTC AGC GCC GAC GAC GGA GAC TAC GCT CAG CAG CTT CCC GAG TAT CCT TCC GTC GCA GCG AAC TCC AGT ACT TCG GAA TCA CAC GAT CAG CAG TCC CTA CTG TTG GTG CCA ACG ATC GCA ACG AAG CTG TCG AAC TTG CTG ACC GAC TTG GTC AGT GTT GGT GGG GGA ACC GGG GGA GTG GCC GGC AGT GTC AAT TTC AGT GCA TTG TTG TCG CTC AAT TTC AGC ACC AGT GGC AAT AGC AGC TCT ATT GGT GTC AGT GCA ACA GCA ACA ACG GCC TTA GCG GTG GCG TTT GAC GAT CAT CTA CCA TCA TCG GAA CGC ACG CAG AAC CAC TCC ACC GAT GGA AGG TTC AAC TGC GAC GGT CGC AAC CTG GCG GAA TTG GCA ACG AGT GCG TTC GGA AGT GCG ACC GGT GGG GGC GCG AGT GGC GAT GGG GGG TTC GGT CCG AAA TTG GAA GTT TCA AAA ACG ATG GAC GTG ATC GGG AAC AGT GGT GGT GTT GGT GAT GGA ACC GAT TCG GTC GGG TTC GGC ATC GGA GAC GGC GGT GCG GCC ACG CAA CTG TTG ATT GCC AAC TGG CAG GAC TGT GTC GTT GTG ATA TTA TTT TGC TTA CTG ATC GTA GTG ACT GTT ATC GGC AAC ACG CTC GTG ATC CTG TCG GTG ATT ACC ACT CGG CGG CTG CGG ACC GTC ACC AAC TGC TTC GTG ATG AGC CTG GCC GTG GCC GAT TGG TTG GTC GGG ATT TTC GTA ATG CCC CCG GCT GTC GCG GTC CAT CTT TTG GGA TCG TGG CCG CTC GGA TGG ATA CTG TGT GAT ATA TGG ATC TCA TTA GAT GTT CTA CTG TGT ACA GCG TCA ATC CTT AGC TTA TGT GCA ATT AGT ATA GAT AGA TAT CTA GCG GTA ACG CAA CCG CTC AAC TAC TCT CGC CGA CGG CGC TCG AAA CGG CTG GCC CTC CTG ATG ATT CTG GTC GTC TGG GTA CTA GCA CTG GCC ATC ACC TGC CCT CCC ATT CTC GGC TGG TAT GAA CCG GGC CGG CGG GAG CTC ACC CAG TGC CGG TAC AAC CAA AAC GAG GGC TAC GTC GTC TTC TCG GCG ATG GGA TCC TTC TTC ATT CCG ATG GCC GTC ATG ATC TAC GTG TAC GTG CGG ATA TCC TGC GTG GTG GCA TCC CGC CAC GAC AAA ATG ACC GAA ATC GAA GTG CAC AAG AAA AGT CAC CGA ACG CGT GAT GCC GAC GAG CCG TGC TAC CAC TAC GCT TCG GAG ATG GAT CCG GCC CAG TTG AGT CCC AAG CGA AAG CGT TCC TCA TCG CAG CTC TCG ACA GCA ACC ACA ACT TAC GTG ACA CAG CTG GGG CCA GGC GGT GGC GGA GGC TCA TCC AGC TTT ACG GCC CTT AGC GGA GAC TGC CCT GGG GGA GAT TGC CTT AAG CTG GGG GGA GGC GGC ACC AGA AAA AAT CAC AAG AAT GGA CAT TAC GAG CTT GTC GAC ATC AAT TCG ATG TCC AAA ACC ACA ACT TCT TTC GTG GGC AAC GGT GGA GGT ACG AAC GGA GCT GGA CTC GAT GAG GAC CAT ATG GAC CCG GGT GGC CAG CGG GAG TCA CCG GGA GCT ACC GGA GGC ATA AAA CGC AAC TCC ACC ATC CAC TAC AAG CTG GCC TCG ACG ACG GAG CTT TGG TCC TCG GCC AAT AAC GTT TTC AAT CAA CAG CAG AAA CCA CAA ACG GCA AAT CAA TCG GGC GGC TTG AGG CGT ACG AAC ACC ATT AGT AAC TCA ATT ACG TCG TCG ACC GCC ACG GCA TCG GCC TCG GCA GTG GCG GCC GGG CGA AAT ATC CGG ATC CAC CAG AAG TCA CTT TCG TCG CGA ATT TTG TCC ATG AAG CGA GAG AAC AAA ACG ACC CAA ACG CTC AGC ATC GTG GTC GGA GGC TTC ATC GCG TGC TGG CTG CCC TTC TTC ATC CAT TAC ATC ATT ACG CCG TTC CTG CCA GAA GAG CTG GCT GCA CCC AAA CTA GGG GAG TTT TTC ACA TGG CTC GGC TGG ATC AAC AGC GCC ATC AAT CCG TTC ATC TAC GCG TTC TAC AGC GTG GAC TTC CGG GCG GCC TTC TGG CGC TTG ACG TTG CGG CGC TTT TTC CGC AAC AGC GAG AAG GCA CCC TTT GCC AAC ATT CAC AAC ATG TCC ATG CGG AGA TAA

METRLDDRCAALLLQLSADDGDYAQQLPEYPSVAANSSTSESHDQQSLLLVPTIATKLSNLLTDLVSVGGGTGGVAGSVNFSALLSLNFSTSGNSSSIGVSATATTALAVAFDDHLPSSERTQNHSTDGRFNCDGRNLAELATSAFGSATGGGASGDGGFGPKLEVSKTMDVIGNSGGVGDGTDSVGFGIGDGGAATQLLIANWQDCVVVILFCLLIVVTVIGNTLVILSVITTRRLRTVTNCFVMSLAVADWLVGIFVMPPAVAVHLLGSWPLGWILCDIWISLDVLLCTASILSLCAISIDRYLAVTQPLNYSRRRRSKRLALLMILVVWVLALAITCPPILGWYEPGRRELTQCRYNQNEGYVVFSAMGSFFIPMAVMIYVYVRISCVVASRHDKMTEIEVHKKSHRTRDADEPCYHYASEMDPAQLSPKRKRSSSQLSTATTTYVTQLGPGGGGGSSSFTALSGDCPGGDCLKLGGGGTRKNHKNGHYELVDINSMSKTTTSFVGNGGGTNGAGLDEDHMDPGGQRESPGATGGIKRNSTIHYKLASTTELWSSANNVFNQQQKPQTANQSGGLRRTNTISNSITSSTATASASAVAAGRNIRIHQKSLSSRILSMKRENKTTQTLSIVVGGFIACWLPFFIHYIITPFLPEELAAPKLGEFFTWLGWINSAINPFIYAFYSVDFRAAFWRLTLRRFFRNSEKAPFANIHNMSMRR*

**Figure S6b.** Nucleotide sequence of the type 2 tyramine receptor (AaTAR2) open reading frame cloned from *Aedes aegypti* and deduced amino acid sequence. The primer used in RT-qPCR were highlighted in red.

**Figure S6c.**

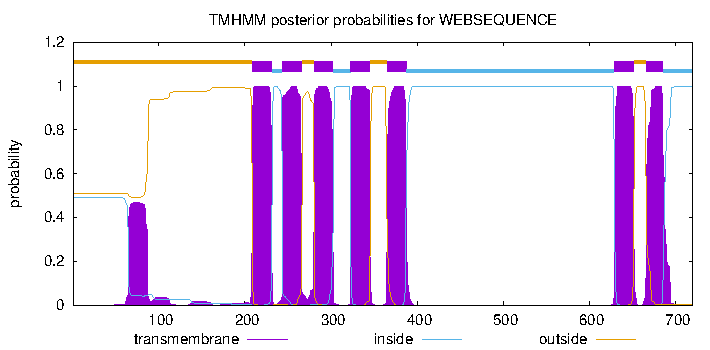

**Figure S6c.** Prediction of the AaTAR2 transmembrane segments obtained with TMHMM v. 2.0 software.

**Figure S6d.**

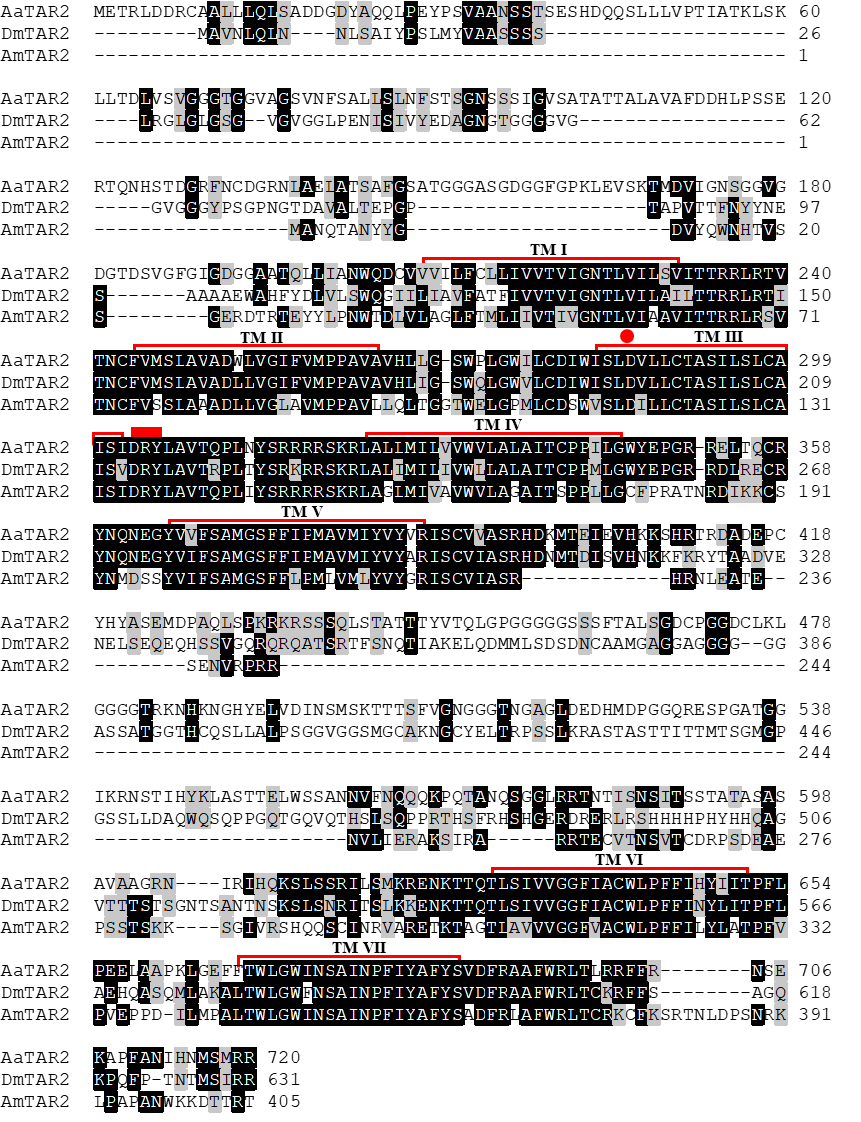

**Figure S6d.** Amino acid sequence alignment of AaTAR2 with orthologous receptors from *D. melanogaster* (DmTAR2) and *A. mellifera* (AmTAR2). The putative seven transmembrane domains (TM I–VII) are indicated by red lines. Identical residues are highlighted in black while conservative substitutions are in grey. A red dot indicates the conserved aspartic acid and the serine residues that could interact with TA.
