## Supplementary material for "Octopamine and tyramine signaling in *Aedes aegypti:* characterization, distribution and potential role in the Dengue vector development and physiology": S7. AaTAR3.docx

**Supplementary figures 7.** *Aedes aegypti* type 3 tyramine receptor (AaTAR3).

**Figure S7a.**

GTCAGTCATTGGTGATTGATCTTACTGGATAATCGAACAGCCACTGTGCTTCCATCTCATCTCCCGGTAAGCCCCATAATT**ATG**GCCAATGAAAGCGGCGCCTTCCCAATCCCCGGCGCCACCACAACGTCAGTGGCATCGATCACCATCAACTACACCAGCACCAGCAGTTCGTTCACCACCAGTCTGTACCAGGAAATCTACGTCAATTTGCAGCTCGTGAACGGCGACCTGCTCAATTTTAGCGAACGTCTCGTAGTGGACCCGAACGGCTCGGCAATGGCGTCCAGCTCCCTGACGGCTTCGTCTTCACCACTTTCTTCCGCATCCCTGATCCCAACATCATCCTTCCACACCGGAATGGTAGCCACGGCTGCGGCAGACACTTCCACCGACAAGGACTGCACCAAGATTTGCCTGTCCTGGGAGAAAATGTTTCTGGTGATTTTGTTCTGTGCGCTGATTGTCATAACGGTGATTGGGAACACTCTGGTGATCCTTTCGGTGGCCACGACCCGACGGTTGAGGACCGTTACCAATTGTTTCGTCATGAGCTTGGCCGTGGCCGATTGGCTTGTGGGAATCTTCGTGATGCCTCCGGCTGTGATAGTTTTTGTTGTAGAGAAGTGGCAACTGGGTTGGATACTCTGTGATATTTGGATATCGCTCGATGTCCTCCTCTGCACGGCCTCGATTCTGAGTCTGTGTGCAATCAGTGTCGATAGATACCTGGCCGTAACTCAACCCCTAACGTACTCAAAACGAAGACGTTCCAAAAGGCTGGCCCTGCTGATGATTTTCGTCGTTTGGCTCGTGGCACTGGCCATCACGTGCCCTCCCATCCTCGGATGGTACGACCAGGACCGGGAGCGGAACGAGTGCCAGTACAACCAAAACAAGGGATACGTGGTTTTTTCGGCGATGGGTTCCTTCTTCATACCGATGAGCGTGATGCTGTACGTCTACTCGAAGATTTGCTGCGTGCTCACGTCCCGGCAGCACCGGATGACCAAGACCGAGGCTACCGAGAAAAACTGCGAGATCGACATCGACAACTATACCTCGGAGATAGACACCAGTCCGAAGCAGGGAACCTACGTGAGCCGTAATATATCGTCCAATTATGCCAACTCGCACCAAACGCTGTACGAGTTCTTGAGTTCGGCTCGCAACACTCCGAACGCCACGCGCCACCAGTTGACAGCACCGAGGTCTTCCGCAGTCGGGCTGACAACGGTCATGGGCGGCGGCAACGGGAACAACAACGGCAATAGCCACTACGAGATGAGGCCAATCAACACGGTCCAGTTCAGTGCCAGCCAGATTTCGCTGGCCGAATCCTGCGTTTCCAACGTGACCAACAGCGGACTAAACGCCACCAAAGTTCGAGTCCATGGCGGCAAACGAATCCCCATACGGATTTCATCGCTGAAGCGCGAAACAAAAACTGCCCAAACTCTGAGCATGGTGGTGGGCGGCTTTATTGCCTGTTGGCTACCCTTCTTCGTGTACTACCTGTTGATGCCGTTCCTGCCGGAGCATTCGCAGAGCAAACCTCTGATGGCATTTCTCACATGGCTCGGTTGGATCAACAGCGCCATCAATCCGTTTATCTACGCATTCTACAACGTGGACTTCCGGATCGCCTTCTGGCGGTTGACGTTTAGGAAGTTCTACAAAAACAAGCACAATTTGGCGTTTTTTAAGTCG**TAG**GATGGAGGGGGTAGGATGTGGTGCAGAAGTGGTGGGGAACGACGACGAGTGCTTACGGTAGACCGTGGAATGTTTTCCGATAACAAAGCAGCTACGGTGACCTTTGTGGAGGAATGTGGCGTTTGAAGATGAGAGATTGAAAGGTACCGGGATTGCGGAGATTTTACATATGAATTTCACATGAGAATACAAATGGGGGCCCCAATAGCCGTAGTTGTAAATAACGCAGCCATTCATCAAGACCAAGCTGTGGCTTGTGGG

**Figure S7a.** *Aedes aegypti* type 3 tyramine receptor (AaTAR3) complete annotated sequence (XM_021837306.1). The primer used to amplify AaTAR3 were highlighted in blue. The start and the stop codon were highlighted in yellow.

**Figure S7b.**

ATG GCC AAT GAA AGC GGC GCC TTC CCA ATC CCG GGC GCC ACC ACG ACG TCA GTG GCA TCG ATC ACC ATC AAC TAC ACC AGC ACC AGC AGT TCG TTC ACC ACC AGT CTG TAC CAG GAG ATC TAC GTC AAT TTG CAG CTC GTG AAC GGC GAC CTG CTC AAT TTT AGC GAA CGT CTC GTA GTG GAC CCG AAC GGC TCG GCA ATG GCG TCC AGC TCC CTG ACG GCT TCG TGT TCA CCA CTT TCT TCC GCA TCC CTG ATC CCA ACA TCA TCC TTC CAC ACC GGA ATG GTA GCC ACG GCT GCG GCA GAC ACT TCC ACC GAC AAG GAC TGC ACC AAG ATT TGC CTG TCC TGG GAG AAG ATG TTT CTG GTG ATT TTG TTC TGT GCG CTG ATT GTC ATA ACG GTG ATT GGG AAC ACT CTG GTG ATC CTT TCG GTG GCC ACG ACC CGA CGG TTG AGG ACC GTA ACC AAT TGT TTC GTC ATG AGC TTG GCC GTG GCC GAT TGG CTT GTG GGA ATC TTC GTG ATG CCT CCG GCT GTG ATA GTT TTT GTT GTA GAA AAG TGG CAA CTG GGT TGG ATA CTC TGT GAC ATT TGG ATA TCG CTC GAT GTC CTC CTC TGC ACG GCC TCG ATT CTG AGT CTG TGT GCA ATC AGT GTC GAT AGA TAC CTG GCC GTA ACT CAA CCC CTA ACG TAC TCA AAA CGA AGA CGT TCC AAA AGG CTG GCC CTG CTG ATG ATT TTC GTC GTT TGG CTC GTG GCA CTG GCC ATC ACG TGC CCT CCC ATC CTC GGA TGG TAC GAC CAG GAC CGG GAG CGG AAC GAG TGC CAG TAC AAC CAA AAC AAG GGA TAC GTG GTA TTT TCG GCG ATG GGT TCC TTC TTC ATA CCG ATG AGC GTG ATG CTG TAC GTC TAC TCG AAG ATT TGC TGC GTG CTC ACG TCC CGG CAG CAC CGG ATG ACC AAG ACC GAG GCT ACC GAG AAA AAC TGC GAG ATC GAC ATC GAC AAC TAT ACC TCG GAG ATA GAC ACC AGT CCG AAG CAG GGA ACC TAC GTG AGC CGT AAT ATT TCG TCC AAT TAT GCT AAC TCG CAC CAA ACG CTG TAC GAG TTC TTG AGT TCG GCT CGC AAC ACT CCG AAC GCC ACG CGC CAC CAG TTG ACA GCA CCG AGG TCT TCC GCA GTC GGG CTG ACA ACG GTC ATG GGC GGC GGC AAC GGG AAC AAC AAC GGC AAT AGC CAC TAC GAG ATG AGG CCA ATC AAC ACG GTC CAG TTC AGT GCC AGC CAG ATT TCG CTG GCT GAA TCC TGC GTT TCC AAC GTG ACC AAC AGC GGA CTA AAC GCC ACC AAA GTT CGA GTC CAT GGC GGC AAA CGA ATC CCC ATA CGG ATT TCA TCG CTG AAG CGC GAA ACA AAA ACT GCC CAA ACT CTG AGC ATG GTG GTG GGC GGC TTT ATT GCC TGT TGG CTA CCC TTC TTC GTG TAC TAC CTG TTG ATG CCG TTC CTG CCG GAG CAT TCG CAG AGC AAA CCT CTG ATG GCA TTT CTC ACA TGG CTC GGT TGG ATC AAC AGT GCC ATC AAC CCG TTT ATC TAC GCA TTC TAC AAC GTG GAC TTC CGG ATC GCC TTC TGG CGG TTG ACG TTT AGG AAG TTC TAC AAA AAC AAG CAC AAT TTG GCG TTT TTT AAG TCG TAG

MANESGAFPIPGATTTSVASITINYTSTSSSFTTSLYQEIYVNLQLVNGDLLNFSERLVVDPNGSAMASSSLTASCSPLSSASLIPTSSFHTGMVATAAADTSTDKDCTKICLSWEKMFLVILFCALIVITVIGNTLVILSVATTRRLRTVTNCFVMSLAVADWLVGIFVMPPAVIVFVVEKWQLGWILCDIWISLDVLLCTASILSLCAISVDRYLAVTQPLTYSKRRRSKRLALLMIFVVWLVALAITCPPILGWYDQDRERNECQYNQNKGYVVFSAMGSFFIPMSVMLYVYSKICCVLTSRQHRMTKTEATEKNCEIDIDNYTSEIDTSPKQGTYVSRNISSNYANSHQTLYEFLSSARNTPNATRHQLTAPRSSAVGLTTVMGGGNGNNNGNSHYEMRPINTVQFSASQISLAESCVSNVTNSGLNATKVRVHGGKRIPIRISSLKRETKTAQTLSMVVGGFIACWLPFFVYYLLMPFLPEHSQSKPLMAFLTWLGWINSAINPFIYAFYNVDFRIAFWRLTFRKFYKNKHNLAFFKS*

**Figure S7b.** Nucleotide sequence of the type 3 tyramine receptor (AaTAR3) open reading frame cloned from *Aedes aegypti* and deduced amino acid sequence. The primer used in RT-qPCR were highlighted in red.

**Figure S7c.**

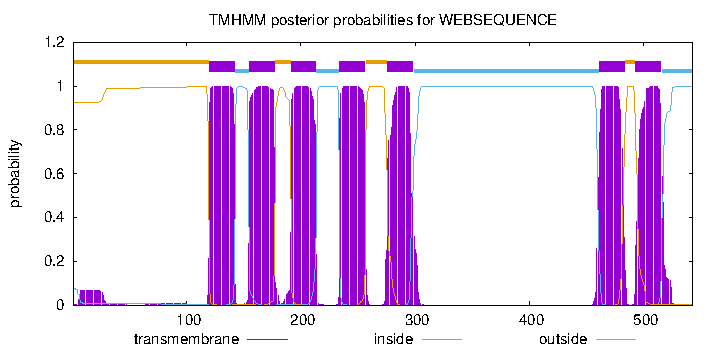

**Figure S7c.** Prediction of the AaTAR3 transmembrane segments obtained with TMHMM v. 2.0 software.

**Figure S7d.**

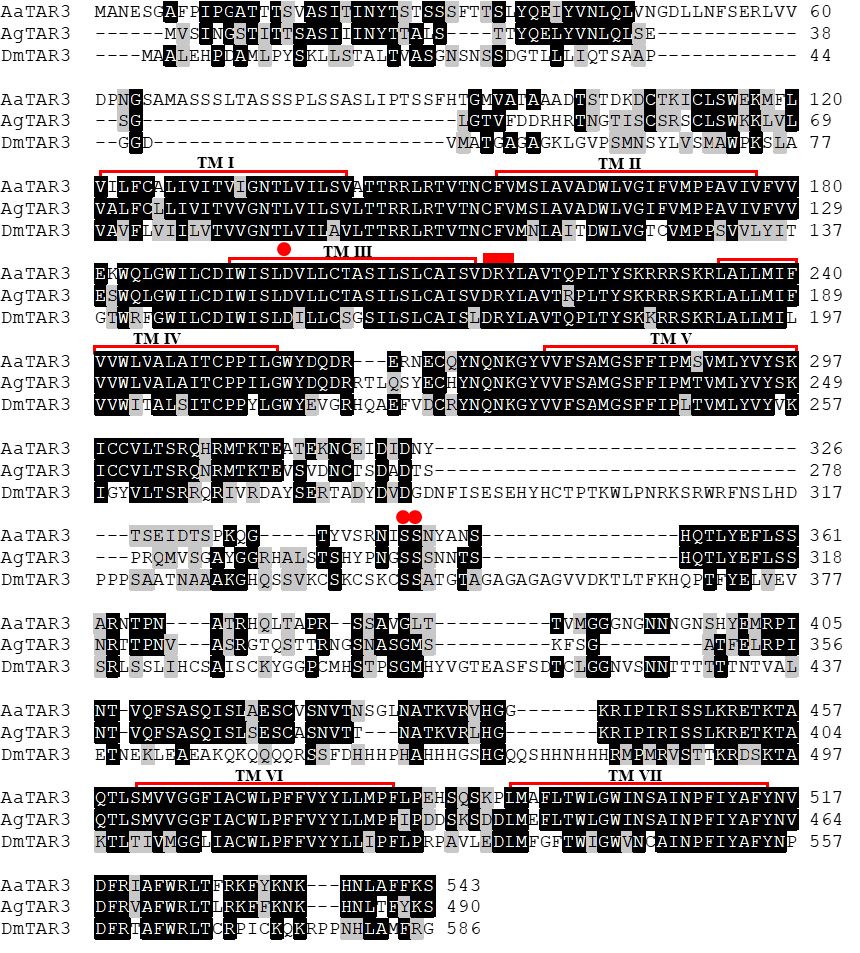

**Figure S7d.** Amino acid sequence alignment of AaTAR3 with orthologous receptors from *A. gambiae* (AgTAR3) and *D. melanogaster* (DmTAR3). The putative seven transmembrane domains (TM I–VII) are indicated by red lines. Identical residues are highlighted in black while conservative substitutions are in grey. A red dot indicates the conserved aspartic acid and the serine residues that could interact with TA.
