## Supplementary material for "Octopamine and tyramine signaling in *Aedes aegypti:* characterization, distribution and potential role in the Dengue vector development and physiology": S8. Feed blood scheme.docx

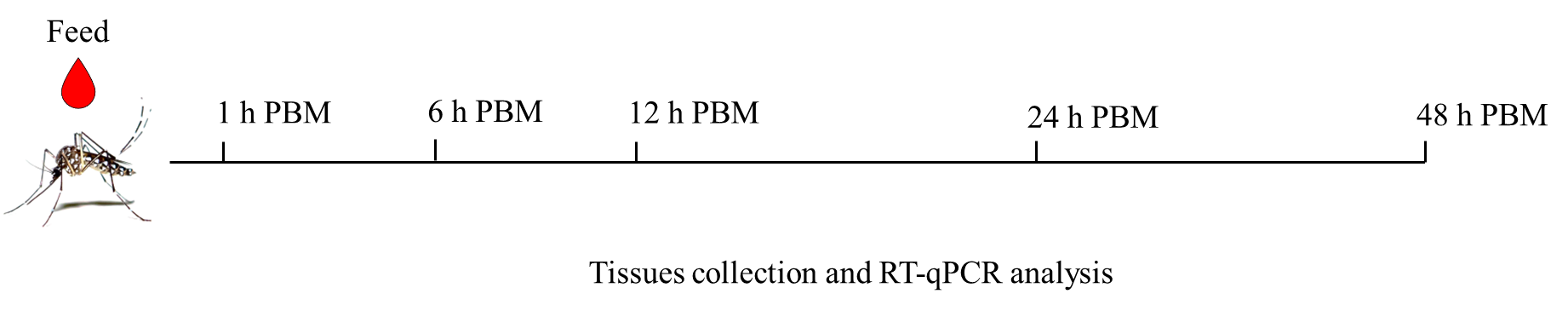

**Figure S8.** Schematic of the protocol used in post blood meal (PBM) experiments. Three days post-ecdysis female *A. aegypti*, 12 hours starved, were fed with blood, and reared under a 12:12 light: dark cycle at 26 °C. The insects were collected at several time points PBM (1, 6, 12, 24, and 48 hours) for tissue/organ collection and subsequent RT-qPCR analysis.
