## Supplementary material for "Octopamine and tyramine signaling in *Aedes aegypti:* characterization, distribution and potential role in the Dengue vector development and physiology": T1. Primer sequences.docx

| Gene | Primer sequence (5’-3’) |
| --- | --- |
| Primers used for OA and TA sequences amplification and sequencing | |
| AaOAα1-R | Fw: TTCTGGTCCGAGCTGAGATT |
|  | Rev: CCTTAAAACCACGCGGAATA |
| AaOAα1-R (ORF) | Fw: **ATG**AATGCGACCGAGTGC |
|  | Rev: **TCA**TCTGCTGTCGGACAGGT |
| AaOAα2-R | Fw: **ATG**GATTACCCAATGATAGCAAC |
|  | Rev: **TCA**CTTGAACAGGATCCGC |
| AaOAβ2-R | Fw: AATTTCGCAAATTTCGTTCG |
|  | Rev: CTGTCACCCGCACAGTTCTA |
| AaOAβ2-R (ORF) | Fw: **ATG**ATGAATCCTTCCAATGACG |
|  | Rev: **TTA**GAGACTCTCGCCGATCTC |
| AaOAβ3-R | Fw: GAGCCAGTCGGTAGTGGAAG |
|  | Rev: TTTTCCAACGACGGATTGAT |
| AaOAβ3-R (ORF) | Fw: **ATG**GCCCTCAGAGCAATGT |
|  | Rev: **CTA**CACGTAGTAGGCGCTGTGT |
| AaTAR1 | Fw: AATCCGTCACAACCAGAAGG |
|  | Rev: ACTCGATTGCTGATGCAGTG |
| AaTAR1 (ORF) | Fw: **ATG**GCAATTGTATCAGTGATACC |
|  | Rev: **CTA**CTGTTTGATTCCGAGCAAT |
| AaTAR2 | Fw: TTGCGGTACTGTTTGCTCTG |
|  | Rev: CATCCGTGTGAAACCACATC |
| AaTAR2 (ORF) | Fw: **ATG**GAAACTCGCCTCGATG |
|  | Rev: **TTA**TCTCCGCATGGACATGT |
| AaTAR3 | Fw: CTGGATAATCGAACAGCCACT |
|  | Rev: AGCCACAGCTTGGTCTTGAT |
| AaTAR3 (ORF) | Fw: **ATG**GCCAATGAAAGCGG |
|  | Rev: **CTA**CGACTTAAAAAACGCCAAAT |
| Primers used for RT-qPCR analysis | |
| AaOAα1-R | Fw: ATCATCGTCGGGTTGTTCAT |
|  | Rev: GGAAAACAGGGCGTAGATCA |
| AaOAα2-R | Fw: GTGGTGAAACCGCTCAAGTT |
|  | Rev: GGTTCCAGTTCGCTACAAGC |
| AaOAβ2-R | Fw: GCAACCTGCTCGTCATCATA |
|  | Rev: CATCCAAACTGTTCCACACG |
| AaOAβ3-R | Fw: GATTCAGGCAGTCCGGTAAA |
|  | Rev: GGGTTGACACTGTCCTCGTT |
| AaTAR1 | Fw: GCATCCACGTATGCAAAATG |
|  | Rev: TGGAGAACTGATCAGCAACG |
| AaTAR2 | Fw: GAGGCTCATCCAGCTTTACG |
|  | Rev: CCCACGAAAGAAGTTGTGGT |
| AaTAR3 | Fw: GTCAATTTGCAGCTCGTGAA |
|  | Rev: GTCCTTGTCGGTGGAAGTGT |
| Actin | Fw: CGTTCGTGACATCAAGGAAA |
|  | Rev: GAACGATGGCTGGAAGAGAG |
| Rps17 | Fw: AAGAAGTGGCCATCATTCCA |
|  | Rev: GGTCTCCGGGTCGACTTC |
| Rps32 | Fw: CAGTCCGATCGCTATGACAA |
|  | Rev: ATCATCAGCACCTCCAGCTC |

**Supplementary table T1.** Primers used in this research work.
