## Supplementary material for "Octopamine and tyramine signaling in *Aedes aegypti:* characterization, distribution and potential role in the Dengue vector development and physiology": T2. Accession numbers.docx

| Accession number | Receptor codified | Abbreviated name | Species |
| --- | --- | --- | --- |
| XP_024667085.1 | Type 3 tyramine receptor | AaTAR3 | *A. gambiae* |
| ACN12797.1 | Type 1 tyramine receptor | AiTAR1 | *A. ipsilon* |
| NP_001011565.1 | Octopamine receptor α1 | AmOAα1 | *A. mellifera* |
| XP_397139.3 | Octopamine receptor β1 | AmOAβ1 | *A. mellifera* |
| XP_006558130.1 | Octopamine receptor β2 | AmOAβ2 | *A. mellifera* |
| XP_006557728.1 | Octopamine receptor β3 | AmOAβ3 | *A. mellifera* |
| NP_001011594.1 | Type 1 tyramine receptor | AmTAR1 | *A. mellifera* |
| NP_001032395.1 | Type 2 tyramine receptor | AmTAR2 | *A. mellifera* |
| PSN51957.1 | Octopamine receptor α 1 | BgOAα1 | *B. germanica* |
| PSN37259.1 | Octopamine receptor α1 | BgOAα2 | *B. germanica* |
| XP_037876464.1 | Octopamine receptor α1 | BmOAα1 | *B. mori* |
| XP_037876565.1 | Octopamine receptor β1 | BmOAβ1 | *B. mori* |
| XP_004928654.1 | Octopamine receptor β2 | BmOAβ2 | *B. mori* |
| NP_001037504.1 | Type 1 tyramine receptor | BmTAR1 | *B. mori* |
| NP_001164649.1 | Type 2 tyramine receptor | BmTAR2 | *B. mori* |
| XP_018897697.1 | Octopamine receptor α1 | BtOAα1 | *B. tabaci* |
| XP_018900354.1 | Octopamine receptor β2 | BtOAβ2 | *B. tabaci* |
| XP_018906763.1 | Octopamine receptor β3 | BtOAβ3 | *B. tabaci* |
| AEQ33589.1 | Octopamine receptor α1 | CsOAα1 | *C. suppressalis* |
| AIC75371.1 | Octopamine receptor α2 | CsOAα2 | *C. suppressalis* |
| AGV79326.1 | Octopamine receptor β1 | CsOAβ1 | *C. suppressalis* |
| AEO89318.1 | Octopamine receptor β2 | CsOAβ2 | *C. suppressalis* |
| AFG26689.1 | Type 1 tyramine receptor | CsTAR1 | *C. suppressalis* |
| ADK91078.1 | Type 2 tyramine receptor | CsTAR2 | *C. suppressalis* |
| NP_732542.1 | Octopamine receptor α1 | DmOAα1 | *D. melanogaster* |
| NP_001262715.1 | Octopamine receptor α2 | DmOAα2 | *D. melanogaster* |
| NP_001262843.1 | Octopamine receptor β1 | DmOAβ1 | *D. melanogaster* |
| NP_001303505.1 | Octopamine receptor β2 | DmOAβ2 | *D. melanogaster* |
| NP_001034046.3 | Octopamine receptor β3 | DmOAβ3 | *D. melanogaster* |
| NP_524235 | Odorant receptor 83b | DmO83b | *D. melanogaster* |
| NP_524419.2 | Type 1 tyramine receptor | DmTAR1 | *D. melanogaster* |
| NP_001287382.1 | Type 2 tyramine receptor | DmTAR2 | *D. melanogaster* |
| NP_650651.1 | Type 3 tyramine receptor | DmTAR3 | *D. melanogaster* |
| QEM34249.1 | Type 1 tyramine receptor | DsTAR1 | *D. suzukii* |
| XP_021194395.1 | Octopamine receptor α1 | HaOAα1 | *H. armigera* |
| XP_021187626.1 | Octopamine receptor β1 | HaOAβ1 | *H. armigera* |
| XP_021187646.1 | Octopamine receptor β3 | HaOAβ3 | *H. armigera* |
| XP_014278335.1 | Type 1 tyramine receptor | HhTAR1 | *H. halys* |
| KNC25005.1 | Octopamine receptor β1 | LcOAβ1 | *L. cuprina* |
| XP_023292810.2 | Octopamine receptor β2 | LcOAβ2 | *L. cuprina* |
| Q25321.1 | Type 1 tyramine receptor | LmTAR1 | *L. migratoria* |
| AAK14402.1 | Type 1 tyramine receptor | MbTAR1 | *M. brassicae* |
| XP_039296445.1 | Octopamine receptor α1 | NlOAα1 | *N. lugens* |
| ATY68968.1 | Octopamine receptor β1 | NlOAβ1 | *N. lugens* |
| XP_039276759.1 | Octopamine receptor β2 | NlOAβ2 | *N. lugens* |
| NP_001317479.1 | Octopamine receptor β3 | NvOAβ3 | *N. vespilloides* |
| AAP93817.1 | Octopamine receptor α1 | PaOAα1 | *P. americana* |
| CAQ48240.1 | Type 1 tyramine receptor | PaTAR1 | *P. americana* |
| BAL72847.1 | Type 1 tyramine receptor | PrTAR1 | *P. regina* |
| XP_048486473.1 | Octopamine receptor α1 | PxOAα1 | *P. xylostella* |
| XP_037967488.2 | Octopamine receptor β1 | PxOAβ1 | *P. xylostella* |
| XP_037967636.1 | Octopamine receptor β2 | PxOAβ2 | *P. xylostella* |
| XP_048486417.1 | Octopamine receptor β3 | PxOAβ3 | *P. xylostella* |
| XP_048485348.1 | Type 1 tyramine receptor | PxTAR1 | *P. xylostella* |
| CAA09335.1 | Type 1 tyramine receptor | RmTAR1 | *R. microplus* |
| XP_024085622.1 | Octopamine receptor α1 | RpOAα1 | *R. prolixus* |
| ATI14906.1 | Octopamine receptor β2 | RpOAβ2 | *R. prolixus* |
| ATI14907.1 | Type 1 tyramine receptor | RpTAR1 | *R. prolixus* |
| QVD39297.1 | Octopamine receptor α1 | SgOAα1 | *S. gregaria* |
| NP_001280520.1 | Octopamine receptor α1 | TcOAα1 | *T. castaneum* |
| NP_001280514.1 | Octopamine receptor β1 | TcOAβ1 | *T. castaneum* |
| NP_001280501.1 | Octopamine receptor β2 | TcOAβ2 | *T. castaneum* |
| XP_015838738.1 | Type 2 tyramine receptor | TcTAR2 | *T. castaneum* |

**Supplementary table 2.** Accession number of OA and TA receptors from different insect species used for the maximum likelihood phylogenetic analysis.
